# ER Stress Impedes Regulated Secretion by Governing Key Exocytotic and Granulogenic Molecular Switches

**DOI:** 10.1101/2023.04.18.537291

**Authors:** Mohima Mukherjee, Chandramouli Mukherjee, Vinayak Ghosh, Aamna Jain, Souren Sadhukhan, Sushma Dagar, Bhavani Shankar Sahu

**Affiliations:** National Brain Research Centre, Manesar, Gurgaon, Haryana 122052, India

**Keywords:** Biological Sciences, Cell Biology, Molecular Biology, Biochemistry

## Abstract

Dense core vesicles (DCVs) and synaptic vesicles (SVs) are specialised secretory vesicles (SSVs) in neurons/neuroendocrine cells harbouring cargo whose abnormal release is associated with pathophysiology. Endoplasmic Reticulum (ER) stress and inter-organellar communication are also associated with disease biology. In pursuit of investigating the cell physiological consequences arising from the crosstalk of a stressed ER and DCVs, ER stress was modelled in PC12 neuroendocrine cells using Thapsigargin (Tg). DCV exocytosis was severely compromised in ER-stressed PC12 cells, reversed by Docosahexaenoic acid (DHA). Experiments with Tunicamycin(Tm), an independent ER stressor, yielded similar results. Concurrently, ER stress caused impaired DCV exocytosis also in INS-1 cells. Molecular analysis revealed blunted SNAP25 expression, potentially attributed to augmented levels of ATF4 (a well-known CREB inhibitor) and its transcriptional regulator CREB (also known to regulate key granulogenic players Chromogranin A, Secretogranin II). Our studies revealed severe defects in DCV exocytosis in ER-stressed cells for the first time, mediated by reduced levels of key ‘exocytotic’ and ‘granulogenic’ switches regulated via the CREB/ATF4/eIF2α axis.

## INTRODUCTION

Disruption of Protein homeostasis and vesicle trafficking is associated with pathophysiological conditions such as Alzheimer’s disease (AD) (Ashraf et al., 2014; Barranco et al., 2021), Parkinson’s disease (PD) (Zou et al., 2021), and Amyotrophic Lateral Sclerosis (ALS) (Cicardi et al., 2021; Gagliardi et al., 2021) to name a few (Sweeney et al., 2017; Wang et al., 2014). Endoplasmic Reticulum is the principal site of protein synthesis, folding, and sorting, which gets compromised under ER stress. An ER-stressed cell employs the Unfolded Protein Response (UPR) as a survival strategy mediated by three pathways: IRE1-α, PERK, and ATF6. They initiate the signalling cascades to regulate transcriptional and translational alterations to enhance protein folding and cell survival against ER stress (Shen et al., 2004). ER, stress leads to IRE1-α activation, causing changes in spliced XBP1 levels. XBP1s regulate the activity of several genes, which ultimately upregulates the expression of chaperones to counteract the ER Stress (Unfolded Protein Response). Similarly, ATF6 activation contributes to UPR and determines cell fate to microautophagy. On the other hand, PERK activation attenuates global translation; upon the sustained stressed condition, it induces apoptosis through the PERK ATF4 CHOP pathway (Hetz, 2012). Dense core vesicles (DCVs) are specialized subcellular organelles in neurons and neuroendocrine cells where peptides, hormones and neurotransmitters are stored. They undergo regulated secretion when an appropriate physiological stimulus is provided. DCVs comprise two major structural compartments, the matrix and the membrane. The matrix includes the chromogranin/secretogranin family’s proteins, out of which Chromogranin A (CGA), Chromogranin B (CGB), and Secretogranin II (SCGII) are the most important as they participate in synthesizing neuropeptides into secretory vesicles. Their functionality also lies in regulating the matrix constituent and its exocytosis (Huttner et al., 1991). The release of cargo to the extracellular space is highly regulated and coupled with membrane depolarization and Ca^2+^ influx, ultimately leading to the docking and releasing of the vesicle (Barclay et al., 2005; Neher and Sakaba, 2008). Neuroendocrine cells also have synaptic-like vesicles (SLVs) similar to neurons packed with neurotransmitters which help in intercellular communication (Shi et al., 1998). The release of hormones, neurotransmitters, peptides, and other molecules of physiological relevance is tightly regulated by V-SNARES (VAMP1) and T-SNARES (SNAP25).

Neuropeptides and neurotransmitters are DCV cargo regulating diverse functions such as sleep, behaviour, cardiovascular regulation, body weight, blood glucose levels and their abnormal levels are associated with various diseases, including neurological disorders (Beal and Martin, 1986; Ganten et al., 1991; McCall, 1990; Teleanu et al., 2022; Yang et al., 2021). Likewise, a stressed ER causes disturbances in cellular homeostasis by altering functions such as protein folding and is associated with several conditions such as HD and diabetes (Yoshida, 2007). It is noteworthy that misfolded proteins, such as mutated alpha-synuclein, are associated with disrupted fusion pore formation during exocytosis (Burré et al., 2010; Sulzer and Edwards, 2019), neurodegeneration (Bennett, 2005), and ER Stress (Colla et al., 2012). This unfolds for potential common crosstalk between ER stress and DCV trafficking. Although DCV biogenesis and release have been pursued for decades, their alterations in response to ER stress and subsequent effects are not yet explored.

To study the association of DCV and SLV exocytosis with ER stress, we chose PC12 cells, a well-known cellular model widely used to study the phenomenon of exocytosis and pharmacologically induced ER stress by Tg treatment as described previously (Lindner et al., 2020; Westerink and Ewing, 2008). Following this, we evaluated the functional status of DCV, which suggested a strongly compromised DCV exocytosis in ER-stressed cells. To test the specificity of the contribution of ER stress in impaired DCV exocytosis, we investigated the functional status of regulated exocytosis in ER-stressed cells supplemented with Docosahexaenoic Acid (DHA), an ER stress attenuator (Begum et al., 2014). To our surprise, impaired regulated secretion was restored to near physiological levels in DHA co-treated cells. Transmission electron microscopy(TEM) revealed that ER stress does not change the Size and Number but alters the spatial localisation of DCVs. Molecular analysis revealed ‘expression switching,’ i.e., altered abundances of key granulogenic proteins and exocytotic signature candidates (CGA, SCGII, and SNAP-25) associated with DCV biogenesis and exocytosis via CREB/ATF4. This was further accompanied by enhanced localization of CGA to the lysosomes. Intriguingly, SLVs displayed a impairment of exocytosis.

Furthermore, similar experiments using Tunicamycin, an independent ER Stressor, caused similar effects on impaired exocytosis of regulated secretion. Finally, we have also tested the functional status of DCVs in Tg-mediated ER-stressed INS-1 (Insulin secreting) cells, which also suggested exocytotic defects of DCVs. In summary, our studies, for the first time, provide evidence that ER stress has detrimental effects on DCV exocytosis, plausibly resulting from cumulative effects of expression switching of SNAP25 and chromogranin potentially modulated by CREB/ATF4/EIF2α axis.

## MATERIALS AND METHODS

### EXPERIMENTAL MODEL AND SUBJECT DETAILS

#### Cell Culture and Treatment

Rat PC12 cells originally generated by Greene & Tischler (Greene and Tischler, 1976) were a gift from Nitish Mahapatra, IIT Madras. Early passage cells (< 8) were used for all experiments. The cells were treated routinely with mycoplasma removal reagent (MP Biomedicals, 30-50044). For routine maintenance, the cells were grown in T25 flasks (Nunc, USA) in DMEM GlutaMAX (GIBCO, 12100046), supplemented with 10% horse serum (GIBCO, 16050114), 5% heat-inactivated fetal bovine serum (Sigma Aldrich, F7524), 25mM HEPES (Sigma Aldrich, H3784) and 1% Antibiotic-Antimycotic (GIBCO, 15240062) in 5% CO_2_ at 37°C in a humidified incubator (Sahu et al., 2012). The insulin-secreting cell line INS-1 (a gift from Professor C. B. Wollheim) was cultured in RPMI-1640 (Sigma Aldrich), supplemented with 10% fetal bovine serum (Sigma Aldrich, F7524), 2 mM L-glutamine (GIBCO GlutaMAX, 35050), 25 mM HEPES (Sigma Aldrich, H3784), 1 mM sodium pyruvate (GIBCO, 11360-070), and 50 μM beta-mercaptoethanol (Sigma Aldrich M3148), in a humidified atmosphere (5% CO_2_, 37°C), according to the previously studied method (Asfari et al., 1992). All treatments were performed in a reduced serum media, and 10nM Tg (Sigma Aldrich, T9033) or 0.05 μg/ml Tm (Cayman Chemical, 11445) was used for 24 hours to induce ER stress in PC12 /INS-1 cells. For rescue experiments, 10µM of DHA (Sigma Aldrich, D2534) was used, and cells were primed with DHA for 24 hours before the induction of ER stress (Begum et al., 2014). Cells grown in reduced serum media were used as a control.

#### Cell Viability Assay

MTT assay was used to assess cytotoxicity to optimize our treatment dose for Tg and Tm (van Meerloo et al., 2011). 10^4^/well PC12 cells plated in a 48-well plate were allowed to settle for 24 hours before starting Tg treatment. Media was then replaced with reduced serum media for 2 hours, after which 6 nM, 10 nM, and 30 nM of Tg were administered to the cells for 24 hours. Next, cells were incubated in 0.5 mg/ml of MTT (Sigma Aldrich, 475989) for 4 hours. Formazan crystals formed were dissolved via gentle trituration in DMSO (Sigma Aldrich, 67-68-5). Optical density (OD) was recorded at 570 nm with a microplate reader (Tecan Spark Multimode Reader). The IC_50_ value was defined as the concentration of Tg exhibiting 50% cell viability.

#### Immunoblotting

After treatment, PC12 cells were lysed using 125 µl of RIPA buffer (10 mM Tris-HCl (Sisco Research Laboratory Pvt. Limited, 37969), 150 mM NaCl (MP Biomedicals, 194848), 1 mM NaVO_4_ (Sigma Aldrich, S-6508), 30 mM Na_4_P_2_O_7_ (Sigma Aldrich, 011K0307), 50 mM NaF (Sigma Aldrich, 7681-49-4), 1% Nonidet P-40 (Sigma Aldrich), 0.1% SDS (Affymetrix USB Products, 151-21-3), 1 mM PMSF (Sigma Aldrich, 52K0052), 1% Triton X-100 (Sigma Aldrich, 9002-93-1), 0.5% C_24_H_39_NaO_4_ (Sigma Aldrich, 145224-92-6), and dissolved protease inhibitor cocktail (Roche Diagnostics, 11714900) in water, pH 7.4) and the protein contents of the cell lysate were quantified using a BCA Protein Assay Kit (TAKARA, T9300A) following the manufacturer’s protocol (Song et al., 2021). The samples containing 10 µg protein were separated using a 10% SDS-polyacrylamide gel and transferred onto nitrocellulose membranes. After blocking with 3% BSA (Sigma Aldrich, A4503), membranes were incubated with primary antibodies against EIF2α (Cell Signalling Technology, 9722s, 1:5000), p-EIF2α (Cell Signalling Technology, 9721s, 1:2000), PDI (Cell Signalling Technology, 3501P, 1:3000), SCGII (Proteintech, 20357-1-AP, 1:1000), CGA (Proteintech, 10529-1-AP, 1:3000), CREB (Cell Signalling Technology, 9197s, 1:4000), p-CREB (Cell Signalling Technology, 9191s, 1:1000), SNAP25 (Synaptic Systems, 111011, 1:2500), ATF4 (Proteintech, 10835-1-AP, 1:5000), VAMP 2 (Synaptic Systems, 104202, 1:2000), STX6 (Synaptic Systems, 110062, 1:5000), SYT1 (Synaptic Systems, 105011, 1:1000), HSP90 (Proteintech, 13171-1-AP, 1:2000), GAPDH (Proteintech, 60004-1-Ig, 1:5000), EF-2 (Santa Cruz Biotechnology, SC-166415, 1:5000) and B-TUB (Proteintech, 10094-1-AP, 1:5000) overnight at 4°C followed by incubation with HRP conjugated goat Anti-mouse (1:10,000) and goat Anti-rabbit (1:10,000) secondary antibodies. The membrane was washed 3X10 minutes after primary and secondary antibody incubation to remove excess antibodies. ECL was prepared using the previously described protocol, and bands were visualized using the Uvitec Mini HD9 gel imaging system (Mruk and Cheng, 2011). Densitometry quantifications were done using ImageLab (6.0.1) (Sahu et al., 2019).

#### Quantitative PCR (qPCR)

Primers for qPCR were ordered from IDT for the following genes: *Gapdh, Caspase3, CgA, ScgII, Snap25, Atf4, Xbp1 spliced, Xbp1 unspliced, Bip, Vamp1, Stx1a and Rab27a*. RNA extraction of the samples was done using a Macherey Nagel Nucleospin RNA isolation kit (740955.50), and RNA yields were estimated using a spectrophotometer (NanoDrop ND-1000). The corresponding cDNAs of the samples were prepared using the iScript Select cDNA synthesis kit (Biorad, 1708891). Primers for qPCRs were used at a final concentration of 0.2 µM, and a final amount of **80** ng cDNA was added. The mRNA expression of target genes was determined using the real-time cycler CFX 96 Maestro (Biorad) following the manufacturer’s protocol of iQ SybrGreen (Biorad, 1725121) (Talbot et al., 2004). 2^-ΔΔCq^ was used to calculate the relative expression of the target gene across various treatments normalized to the housekeeping control Gapdh (Schmittgen and Livak, 2008).

*Atf4*: **FP:** CCTGAACAGCGAAGTGTTGG/ **RP: T**GGAGAACCCATGAGGTTTCAA

*Atf6*: **FP:** TCGCCTTTTAGTCCGGTTCTT/ **RP**: GGCTCCATAGGTCTGACTCC

*Caspase3*: **FP:** GGACAGCAGTTACAAAATGGATT/ **RP:** CGGCAGGCCTGAATGATGAAG

*Snap25*: **FP:** CGAAGAGAGTAAAGATGCTGGC/ **RP:** GTTTTGTTGGAGTCAGCCTTCT

*Cga*: **FP:** CGGCAGCATCCAGTTCTCA/ **RP:** AGCCCCTGTCTTTCCATTCA

*ScgII*: **FP:** CCTACTTGAGAAGGAATTTGC/ **RP:** ACCAACCCATTTGGTTTCTC

*Xbp-1 Spliced*: **FP:** TCAGACTACGTGCGCCTCT/ **RP:** TCAGACTACGTGCGCCTCT

*Xbp-1 Unspliced*: **FP:** CTGAGTCCGCAGCAGGTG/ **RP:** CCACATCCGCCGTAAAAGAATG

*Gapdh***: FP:** CGTATTGGGCGCCTGGTCAC/ **RP:** CGGCCTCACCCCATTTGATG

*Bip:* **FP:** GGTACATTTGATCTGACTG/ **RP:** CACTTCCATAGAGTTTGCTG

*Vamp1*: **FP:** GGTTTCCATTGTGTCTGTC/ **RP:** ATCTGTCACATGCCTTTGGT

*Syntaxin 1a*: **FP:** TACAACGCCACTCAGTCAGA/ **RP:** GAGTCCATGATGATCCCAGA

*Rab 27a*: **FP:** TTCAGGGACGCTATGGGTTT/ **RP:** CCGCAGAGCACTATATCTGGG

18s *rRNA:* **FP:** GTAACCCGTTGAACCCCATT/ **RP:** CCATCCAATCGGTAGTAGCG

#### Transfections

PC12 and INS-1 cells were transfected using the manufacturer’s protocol using METAFECTENE® PRO (Biontex, Lot. No. RKP205/RK081621) (Grindheim et al., 2014). In brief, plasmid DNA and METAFECTENE® PRO were mixed in DMEM/RPMI medium without any supplements and incubated for 20 minutes at room temperature to form complexes. These complexes were directly added to the cells. The following plasmids were used: NPY-mApple (Addgene, 83498), NPY-pHTomato (Addgene, 83501), NPY-pHluorin (A kind gift from Sebastian Barg, Uppsala), Synaptophysin-pHTomato (A kind gift from Yulong Li lab, China) and mCherry-Lysosomes-20 (Addgene, 55073). Cells were cultured in G418 containing media for 14 days for stable cell line generation.

#### NPY-mApple Plate Reader Assay

Stably selected NPY-mApple PC12 cells were plated at 3X10^5^ cells/well in a 6-well plate. After that, cells were washed with ice-cold 1X PBS (GIBCO, 21600-069) and then incubated in Tyrode’s buffer (basal: 150 mM NaCl, 5 mM KCl, 2 mM CaCl2, 10 mM HEPES; pH 7.4), (stimulation: 55 mM NaCl, 100 mM KCl, 2 mM CaCl2, 10 mM HEPES; pH 7.4) for 20 minutes. The supernatant was collected, and cells were lysed on ice. Lysed cells were sonicated and spun at 14,000 g for 15 minutes at 4°C. 10 μl of the cellular lysate was used, and the supernatant was diluted at a 1:10 ratio in 1X PBS (GIBCO, 21600-069). 100 µl of the supernatant and diluted cellular lysate was loaded in a dark 96-well plate to eliminate cross-contamination of the fluorescent signal from adjacent wells. NPY-mApple fluorescent signals were then recorded using the TECAN plate reader (Ex λ-568 nm, Em λ-592 nm), subtracting them from the autofluorescence of the plate itself (Wemhöner et al., 2006).

#### Immunofluorescence and Confocal Imaging

3*10^4^ cells/coverslip were seeded onto the PLL-coated coverslips. After treatment, cells were gently rinsed with 1X PBS (GIBCO, 21600-069) and fixed with 4% PFA (FISHER, 30525-89-4) at room temperature for 20 mins, followed by 3X5 minutes wash with 1X PBS to remove excess PFA (FISHER, 30525-89-4). Cells were then permeabilized with 0.1% Triton X-100 (Sigma Aldrich, 9002-93-1) for 10 mins and blocked using 1% BSA (Sigma Aldrich, A4503) for an hour. Further, they were incubated with Goat CgA primary antibody (Santa Cruz Biotechnology, SC-1488, 1:500 in blocking buffer) in a humidified chamber for 1 hr, followed by a gentle wash with 1X PBS to remove the excess antibody. Alexa-fluor 488 conjugated donkey Anti-Goat (Invitrogen, A11055, 1:2000) secondary antibody was then added to the cells for 45 mins in a humidified chamber in the dark. Samples were again rinsed with 1X PBS, and coverslips were mounted using a FlouroshieldTM with dapi (Sigma Aldrich, F6057) to reduce photobleaching (Berlier et al., 2003). Images were captured using a Nikon A1HD25 confocal microscope with 60X oil objective (NA 1.4) at a 512 * 512 pixels resolution. (Representative images were captured at 2048 * 2048 pixels resolution).

#### Live-cell Imaging and Analysis

PC12 cells were transfected with NPY-pHTomato (Addgene) and stably selected, whereas Synaptophysin-pHTomato (Addgene) and NPY-pHluorin (Addgene) were transiently transfected in PC12 and INS-1 cells respectively (Gandasi et al., 2015; Hummer et al., 2017). For imaging, cells were plated on 15 mm coverslips (Assistent/ Glaswarenfabrik, 41001115) coated with 1X Poly-L-Lysine (Sigma Aldrich, P2636), and the same treatment protocol was followed. On the day of the experiment, cells were rinsed with 2.5 mM KCl (MERCK, 61753305001046) containing Tyrode’s buffer (basal), and coverslips were placed in an open imaging chamber (Life Cell Instruments, Korea). Images were captured using Zeiss spinning disk confocal microscope with a Yokogawa CSU-XA1 head at 63X oil (1.4 NA) at 300 ms for 12 sec in the Tyrode’s buffer (basal). At the end of the experiments, Tyrode’s solution containing 100 mM NH_4_Cl, pH 7.4, was added to its final concentration, and images were taken to acquire the total fluorescence of the transfected cell. Captured time-lapse movies were analyzed using ImageJ software, and after background subtraction, the change in puncta intensity/cell (Δf) after 100 mM KCl stimulation as compared to basal fluorescence (f0) was plotted as Δf/f0.

#### Constitutive Secretion

To check the constitutive release of proteins in basal conditions, secreted fractions from unstimulated cells were analyzed by SimplyBlue^TM^ SafeStain (Invitrogen, LC6060) Coomassie staining (Huang et al., 2001; Hummer et al., 2017). PC12 cells were seeded in 60 mm culture dishes at a density of 10^6^ per dish. After 24 hours, treatments were given, followed by media removal and incubation in the Tyrode’s buffer (basal) for 20 mins. The supernatants and cell lysates were collected and resolved using a 10% SDS-polyacrylamide gel. The gel was washed with Milli-Q water twice, then stained with SimplyBlue^TM^ SafeStain (Invitrogen) for 1 hour at room temperature with gentle shaking. To remove the background, the gel was washed with Milli-Q water twice and later imaged in the Azure c300 gel imaging system, and all the automatically detected bands were quantified using ImageLab (6.0.1).

For the HSP-90 secretion experiment, cells were seeded in 6 well plates, and after completion of treatment, cells were incubated with Tyrode’s basal buffer for 3 hours. After that, the supernatant containing basal buffer was collected and resolved in 10% SDS-polyacrylamide gel, following the standard immunoblotting procedure and probed for HSP-90 (Proteintech, 13171-1-AP). Bands were visualized using the Uvitec Mini HD9 gel imaging system (Mruk and Cheng, 2011)

#### Electron Microscopy

Cells were seeded in 100mm cell culture plates with a 60% confluency, and at the end of the treatment, cells were fixed in 2.5% glutaraldehyde (Sigma Aldrich, MKBG0637V) and 2% PFA in 0.15 M Sodium cacodylate buffer (Sigma Aldrich, C0250) and post-fixed in 1% Osmium Tetroxide (Sigma Aldrich, 05500) in 0.1 M Sodium cacodylate buffer for 1h on ice. The cells were then dehydrated in graded series of ethanol (MERCK, 1.00983.0511) (50–100%) on ice, followed by one wash with 100% ethanol and two washes with Propylene Oxide (TCI, E0016) (15 min each) and embedded with Araldite-502 (Ted Pella, 18060). Ultrathin (50 ∼ 60 nm) sections were cut on a Leica UCT ultramicrotome and picked up on carbon-coated copper grids. Sections were stained with 10% Uranyl acetate (Sisco Research Laboratory Pvt. Limited, 81405) for 5-min and Sato’s lead citrate (Electron Microscopy Sciences, 17800) stain for 1-min. Grids were viewed using a JEOL JEM1400-plus TEM (Sahu et al., 2019). Images obtained were measured using Image J software, Dense core vesicle/Dense Core diameters were calculated, and total DCV distribution was obtained by measuring the shortest distance from the cell membrane.

#### Statistical Analysis

Statistical comparisons between two groups were performed with Student’s unpaired t-test when data were normally distributed with similar variances. Otherwise, the Mann-Whitney test was performed to draw the significance. For comparison among multiple groups, One-way ANOVA was performed. A p-value of 0.05 or less was considered statistically significant. Statistical analysis was performed using GraphPad Prism 8.0. The data are presented as means, and error bars show the standard error of the mean (SEM).

## LEAD CONTACT AND MATERIALS AVAILABILITY

Further information and request for resources, reagents and data should be directed to and will be fulfilled by the lead contact, Dr Bhavani Shankar Sahu. This study did not generate new unique reagents.

## RESULTS

### Tg causes ER Stress in PC12 cells without inducing Apoptosis

Thapsigargin (Tg) was used as a pharmacological ER stress inducer in the PC12 cell lines (Szegezdi et al., 2008). We carried out a cell viability assay for various doses of Tg and a specific dose of 10 nM with minimum cell death but able to activate the ER stress pathways were chosen for all subsequent experiments (Fig. 1B) (Brodnanova et al., 2021). At this specific dose, 80% of the cells were viable without enhanced expression of proapoptotic marker Caspase3 (Fig. 1C) but with the activation of the PERK/EIF2α arm of the ER stress pathway, as evident by the activation of p-eIF2α (Fig. 1D). However, the other two arms of the UPR (*Atf6* and *Xbp1*) are unaltered (Supp. Fig. 1A, 1B) in Tg-treated cells. (Shen et al., 2004).

**Figure 1:**
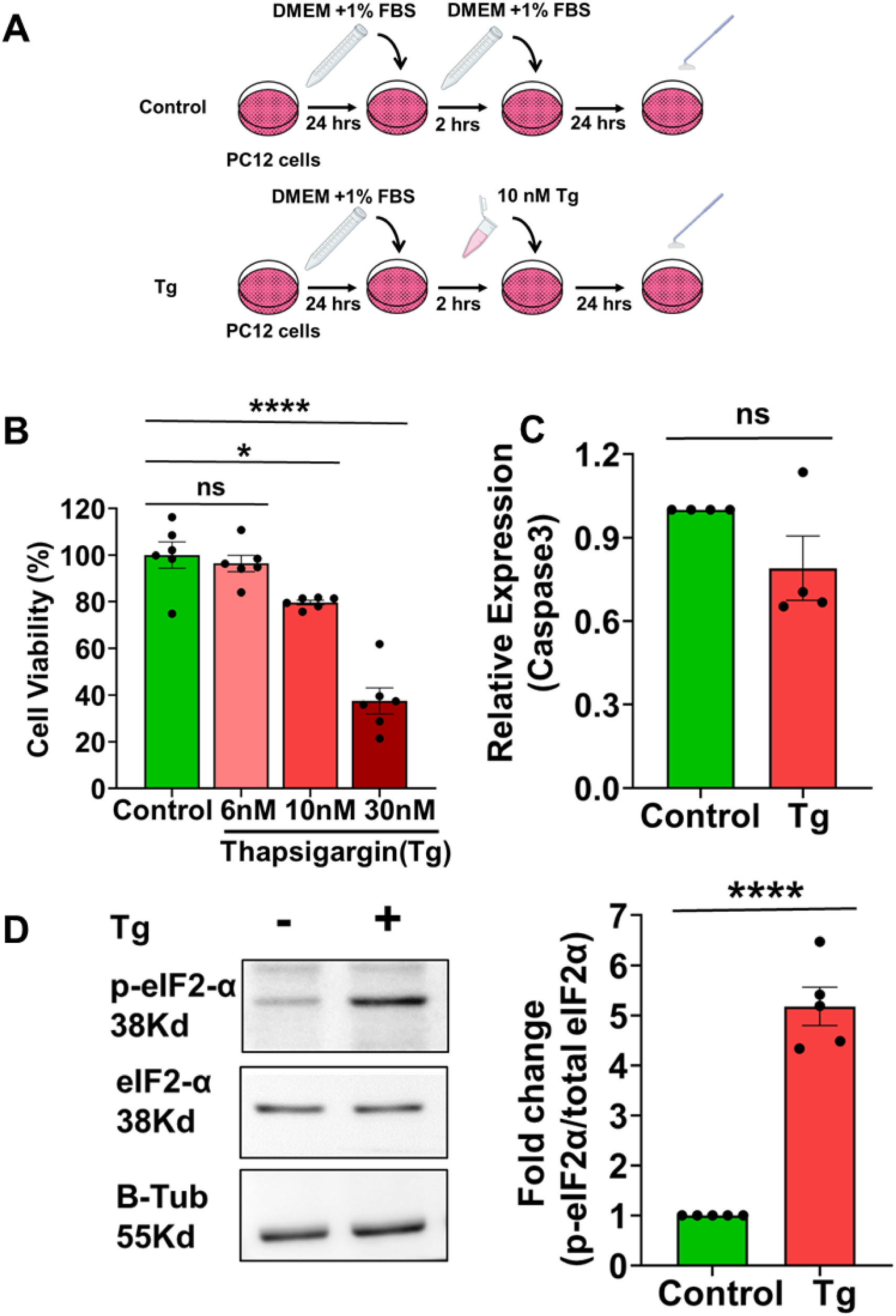
Thapsigargin causes ER Stress in PC12 cells. **A** Treatment protocol: 24 hours after plating, cells were treated with 10 nM Tg under reduced serum conditions, and experiments were performed after 24 hours. **B** Cell viability assay for different Tg doses showed around 80% cell viability in 10nM dosage (ANOVA, N=6). **C** Transcript levels of Caspase 3 in Control and Tg treated cells (Mann-Whitney test, N=4). **D** Representative immunoblot showing p-EIF2α and t-EIF2α (left) expression and its corresponding Quantification (right) in Control and cells treated with Tg. Beta tubulin was used as a loading control (Student’s t-test, N=5). (N denotes the number of Biological replicates). Data are shown as mean ± SEM. (*P < 0.05; **P < 0.01; ***P<0.001; n.s. statistically non-significant).

### ER, stress inhibits exocytosis of dense-core vesicles

NPY-mApple (fluorophore tagged with Neuropeptide Y, a soluble DCV cargo) stable cell lines were generated, and the increase in NPY-mApple fluorescence intensities upon giving an external stimulus was quantified as a measure of regulated secretion. Control cells showed KCl-mediated stimulus-coupled secretion of NPY-mApple, as evidenced by the significantly increased fluorescence in the supernatants and reduced fluorescence intensities of NPY-mApple in the cell lysates compared to their basal unstimulated counterparts. However, Tg-treated ER-stressed cells showed a lack of response to stimulus-coupled secretion accompanied by non-significant changes in NPY-mApple fluorescence, suggesting an impediment in the release of cargo from dense-core vesicles (DCVs) (Fig 2A; Supp. Fig 3C). We also did not observe any change in the global constitutive release signature between Control and Tg-treated cells (Huang et al., 2001; Hummer et al., 2017). (Supp. Fig. 3A, 3B).

**Figure 2:**
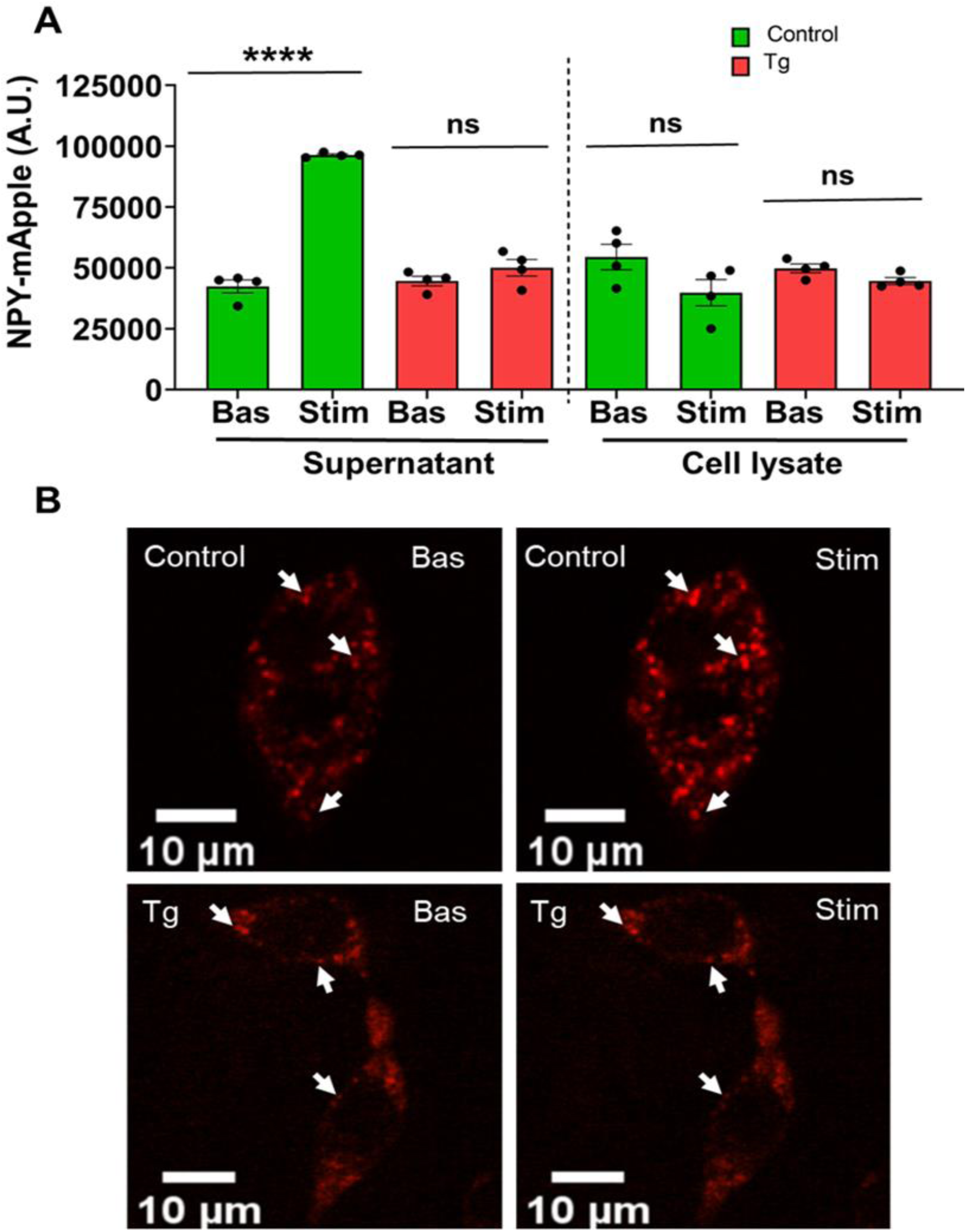
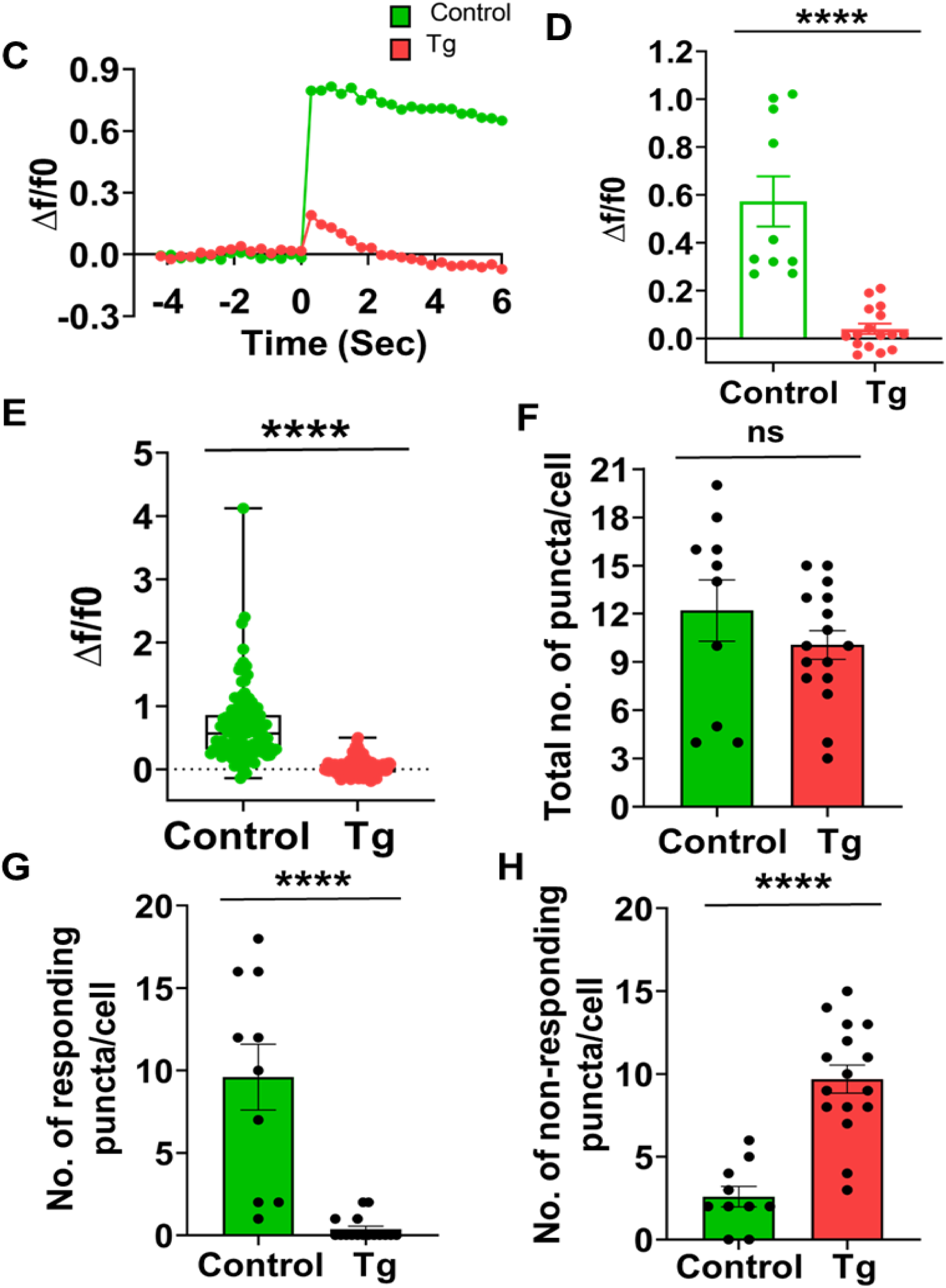
ER stress causes an impediment to regulated secretion in DCVs. **A** Quantitative measure of NPY-mApple fluorescence intensity in Control and Tg-treated PC12 cells transfected with NPY-mApple showing basal and stimulated conditions, measured using a plate reader (Student’s t-test, N=4). **B** Representative images of NPY-pHTomato (red) transfected PC12 cells (Top, Control; bottom, Tg) showing basal and KCl stimulated conditions (white arrows) **C** Representative time traces of mean NPY-pHTomato fluorescence intensities showing an average of all puncta per cell (green, control; red, Tg). **D** Quantitative analysis of the maximum Δf/f0 in control and Tg-treated PC12 cells (Students’t-test, Control, n=10; Tg, n=16). **E** Quantitative analysis showing the maximum Δf/f0 of all NPY-pHTomato puncta from all cells (Mann-Whitney, Control, x=122; Tg, x=161). **(F-H)** Quantification of the total number of NPY-pHTomato puncta/cell (Student’s t-test) **(F)**, No. of responding NPY-pHTomato puncta/cell (Mann-Whitney test) **(G)** No. of non-responding NPY-pHTomato puncta/cell (Student’s t-test) **(H)** considering a minimum of Δf/f0 of 0.3 upon KCl stimulation as a response, (Control, n=10; Tg, n=16) (N=3). (‘N’ denotes the number of biological replicates, and ‘n’ is the number of cells taken for quantification, x= no of puncta analysed) Data are shown as mean ± SEM. (*P < 0.05; **P < 0.01; ***P<0.001; n.s. statistically non-significant).

To further assess the possible defect in DCV exocytosis on Tg treatment, we transfected PC12 cells with NPY-pHTomato (pH-sensitive DCV marker) (Gandasi et al., 2015; Hummer et al., 2017). We then observed the change in NPY-pHTomato puncta fluorescence in response to 100 mM KCl stimulation, and individual exocytotic events were monitored by live-cell imaging using a spinning disk confocal microscope. 100 mM KCl acts as a secretagogue and is known to induce depolarization of plasma membrane leading to the DCVs fusion and exocytosis. We observed increased NPY-pHTomato fluorescence upon 100 mM KCl stimulation in control cells. However, we saw distinct non-responsive dynamics in ER-stressed cells (Fig. 2B, 2C, Supp video 1). Next, we quantified the change in NPY-pHTomato puncta fluorescence (maximum Δf/f0) as a measure of dense core vesicle exocytosis and found a significant reduction in DCV exocytosis in Tg-treated cells as compared to control cells. These findings were in line with the NPY-mApple plate reader-based secretion experiments. Furthermore, we assessed the global response of NPY-pHTomato by analyzing individual exocytotic events. We observed an overall decrease in responsiveness to the stimulus measured as maximum Δf/f0 (Fig. 2D, 2G) and an increase in the number of non-responsive NPY-pHTomato puncta in ER-stressed cells and distribution of all NPY-pHTomato puncta showing maximum Δf/f0 from all cells (Fig. 2E, 2H) (Hummer et al., 2017). Using a similar approach, we observed severe impediments in NPY-pHTomato exocytosis upon secretagogue stimulation in Tg-stressed cells. We have also investigated the functional status of DCV exocytosis in ER-stressed INS-1 cells by NPY-pHluorin imaging and observed a compromised DCV exocytosis (Supp. Figure 2)(A-G), Supp. Video 5) (Ji et al., 2017). Our findings showed an impediment in DCV exocytosis in multiple cellular models of ER stress using multiple ER stressors.

### ER, stress causes defects in regulated exocytosis in SLVs

Neuroendocrine cells also have synaptic-like vesicles (SLVs) apart from DCVs. Extending our observation to SLVs, we investigated the functional status of regulated secretion in SLVs. PC12 cells were transfected with Synaptophysin-pHTomato. Synaptophysin is a membrane glycoprotein localized to synaptic vesicles (SVs) in neurons and SLVs in neuroendocrine cells and hence serves as an important marker for studying the dynamics of such vesicles (Kwon and Chapman, 2011). Individual exocytotic events were tracked by live-cell imaging using a spinning disk confocal microscope under a 63X oil objective. Tg-treated cells showed exocytotic defects and non-responsive dynamics of synaptic-like vesicles upon 100 mM KCl stimulation compared to control cells, which responded with a fluorescence burst upon stimulation with the same secretagogue. (Fig. 3A, 3B, Supp. video. 2). We calculated maximum Δf/f0 as a measure of synaptic vesicle exocytosis, thus reflecting the change in fluorescence between the basal and stimulated conditions. Stressed cells showed reduced to almost absence of response upon stimulation with KCl secretagogue as compared to the control cells (Fig. 3C, 3F) and an increase in the number of non-responsive Synaptophysin-pHTomato puncta in stressed cells (Fig. 3G) and distribution of all Synaptophysin-pHTomato puncta showing maximum Δf/f0 from all cells (Fig. 3D). Hence, a stress-induced impediment to regulated secretion is replicated in SLVs similar to the DCVs.

**Figure 3:**
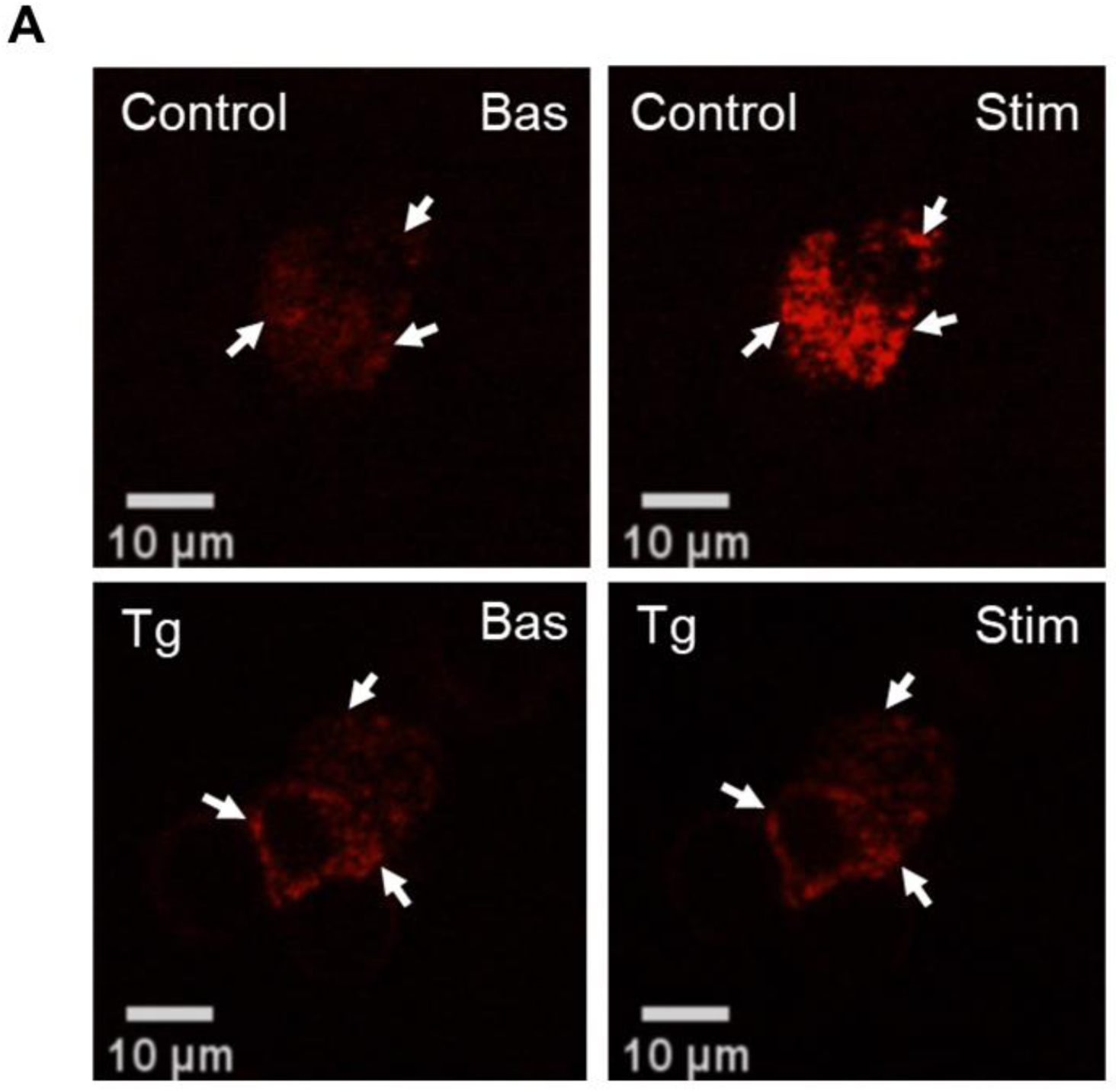
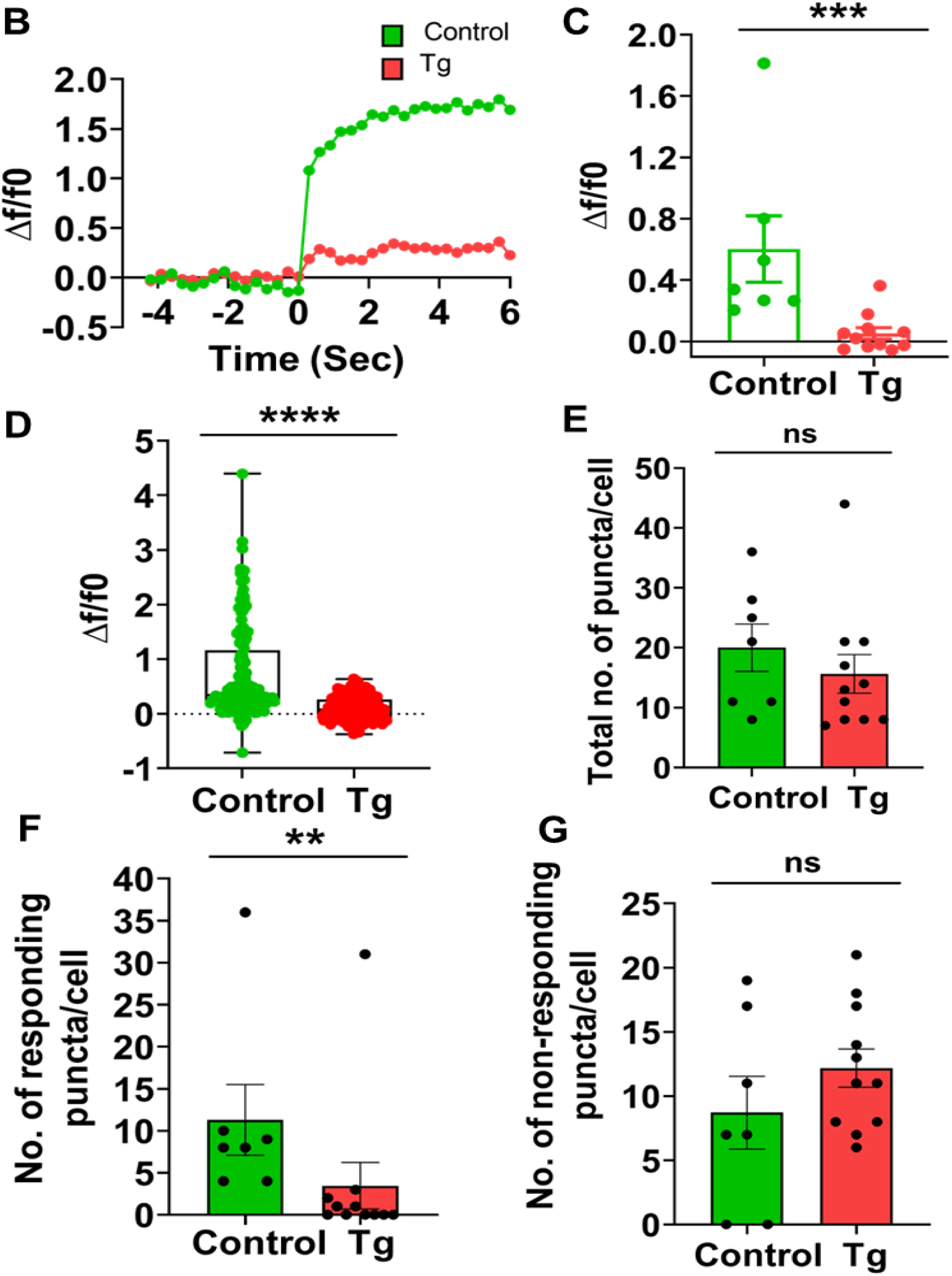
ER stress causes an impediment to regulated secretion in SLVs. **A** Representative image of Synaptophysin-pHTomato (red) transfected PC12 cells (Top, Control; bottom, Tg) showing basal and KCl stimulated conditions (white arrows). **B** Representative time traces of mean Synaptophysin-pHTomato fluorescence intensities showing the average of all puncta per cell (green, control; red, Tg treated cells). **C** Quantitative analysis of the maximum Δf/f0 in control and Tg-treated PC12 cells (Mann-Whitney test, Control, n=7; Tg, n=11, N=3). **D** Quantitative analysis showing the maximum Δf/f0 of all Synaptophysin-pHTomato puncta from all cells (Mann-Whitney test, Control, x=140; Tg, x=172, N=3). Data are shown as mean ± SEM. **(E-G)** Quantification of a total number of Synaptophysin pHTomato puncta/cell (Mann-Whitney test) **(E)** No. of responding Synaptophysin pHTomato puncta/cell (Mann-Whitney test) **(F)** No. of non-responding Synaptophysin pHTomato puncta/cell (Student’s t-test) **(G)** considering a minimum of Δf/f0 of 0.3 upon KCl stimulation as a response, (Control, n=7; Tg, n=11, N=3). (‘N’ denotes the number of biological replicates, and ‘n’ is the number of cells taken for quantification; x= no.of puncta analyzed) Data are shown as mean ± SEM. (*P < 0.05; **P < 0.01; ***P<0.001; n.s. statistically non-significant).

### DHA reverses the defective DCV exocytosis phenotype by reducing ER stress

Docosahexaenoic acid (DHA), one of the most abundant omega-3 fatty acids in the brain, has been previously shown to attenuate ER stress in several in-vitro studies (Begum et al., 2012a, 2013, 2014; Dyall, 2015). In line with previous studies, we see a reduced activation of ER stress as indexed by restoring p-EIF2α/t-EIF2α to near control levels (Fig. 4B). This gave us the first line of data implying the reversal of ER stress phenotype in Tg-treated cells upon DHA co-treatment.

**Figure 4:**
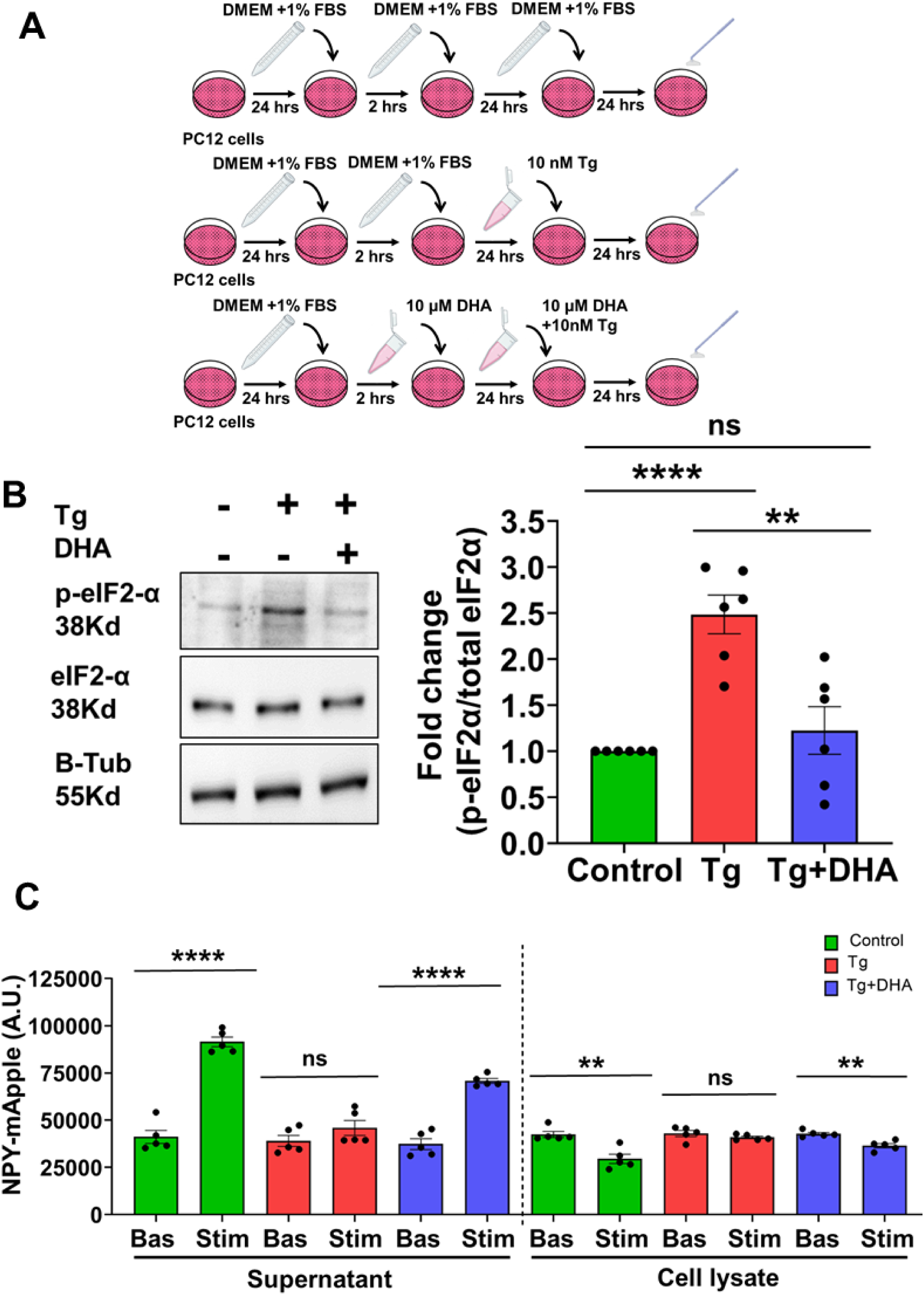
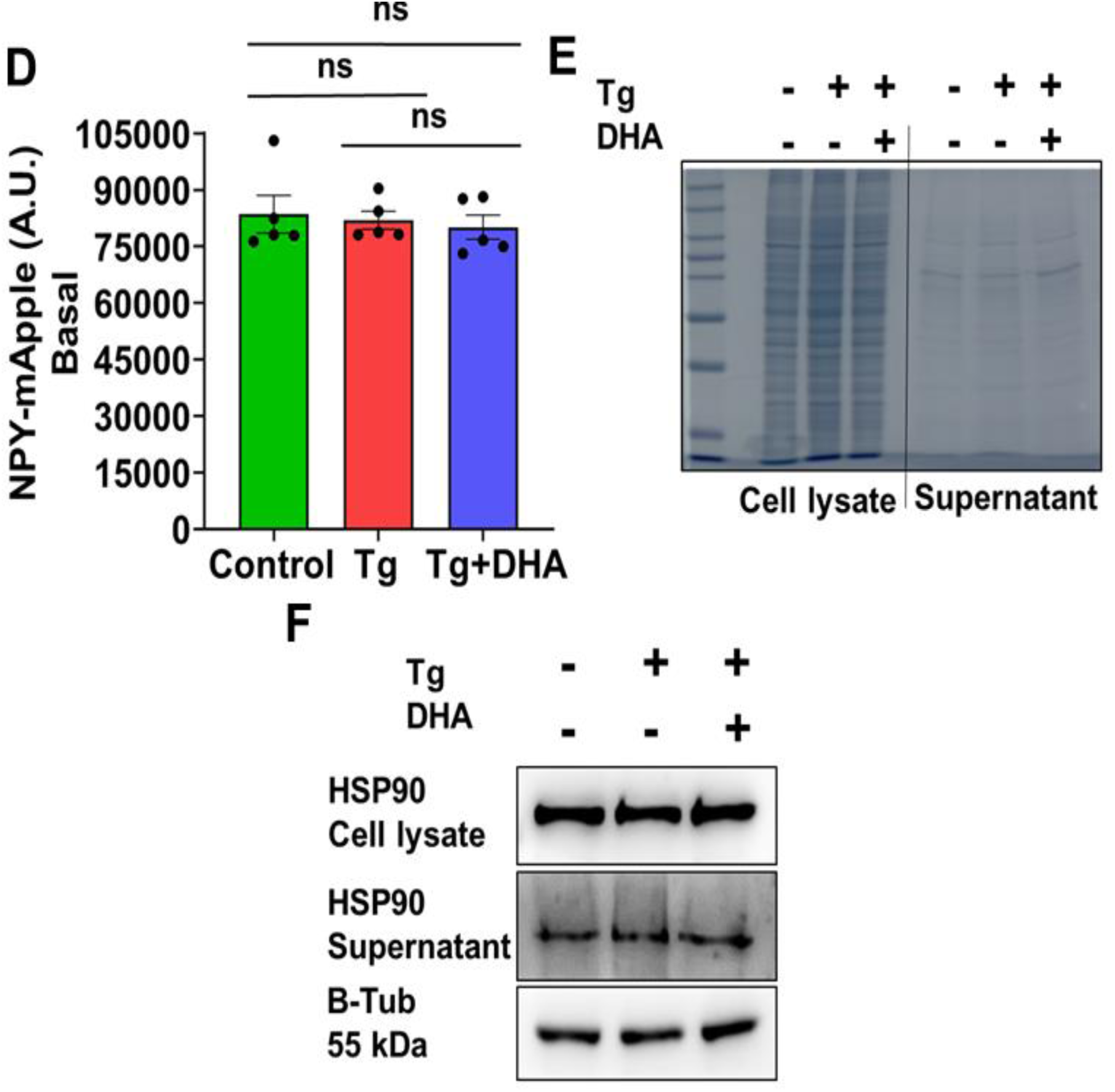
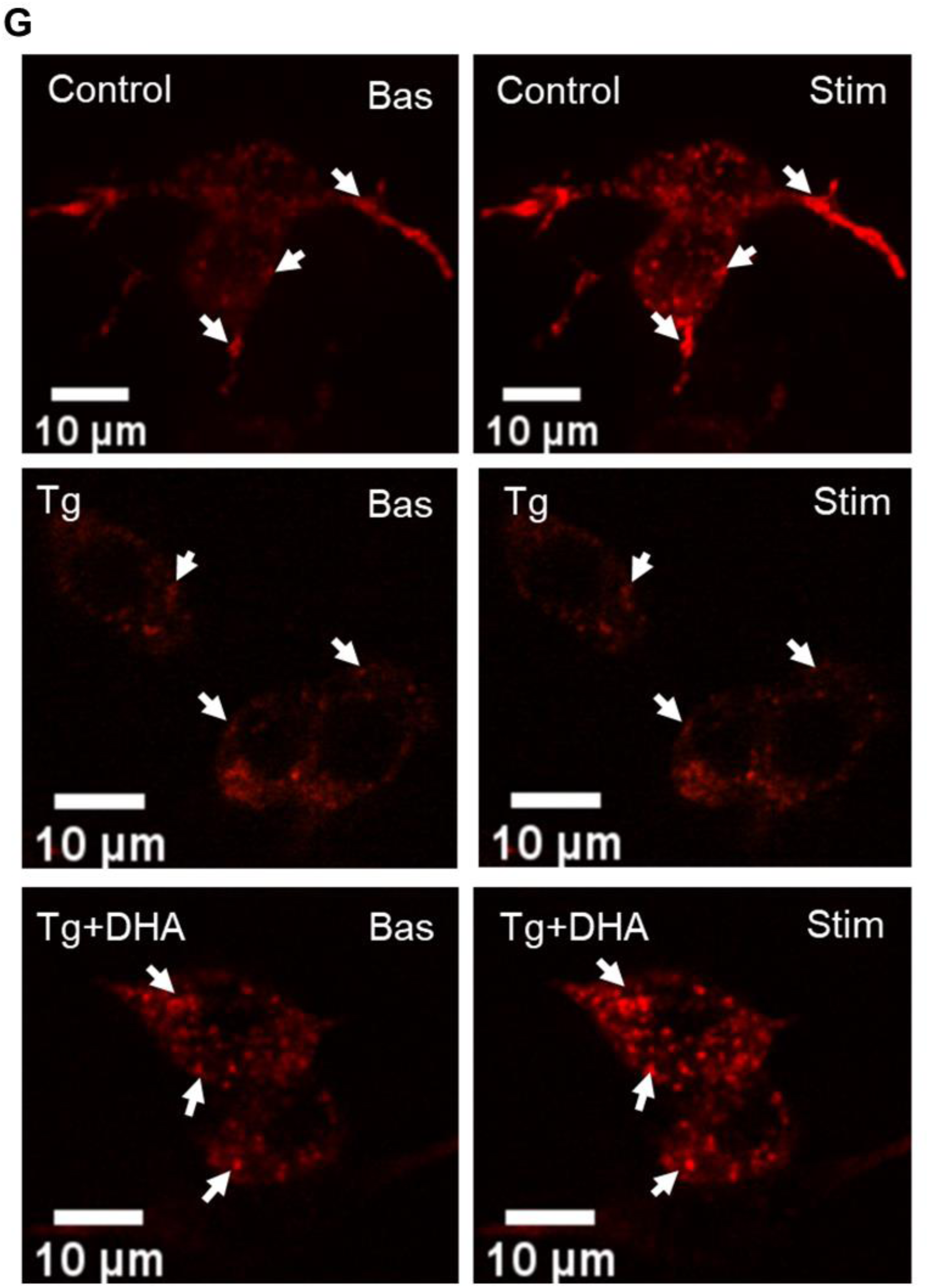
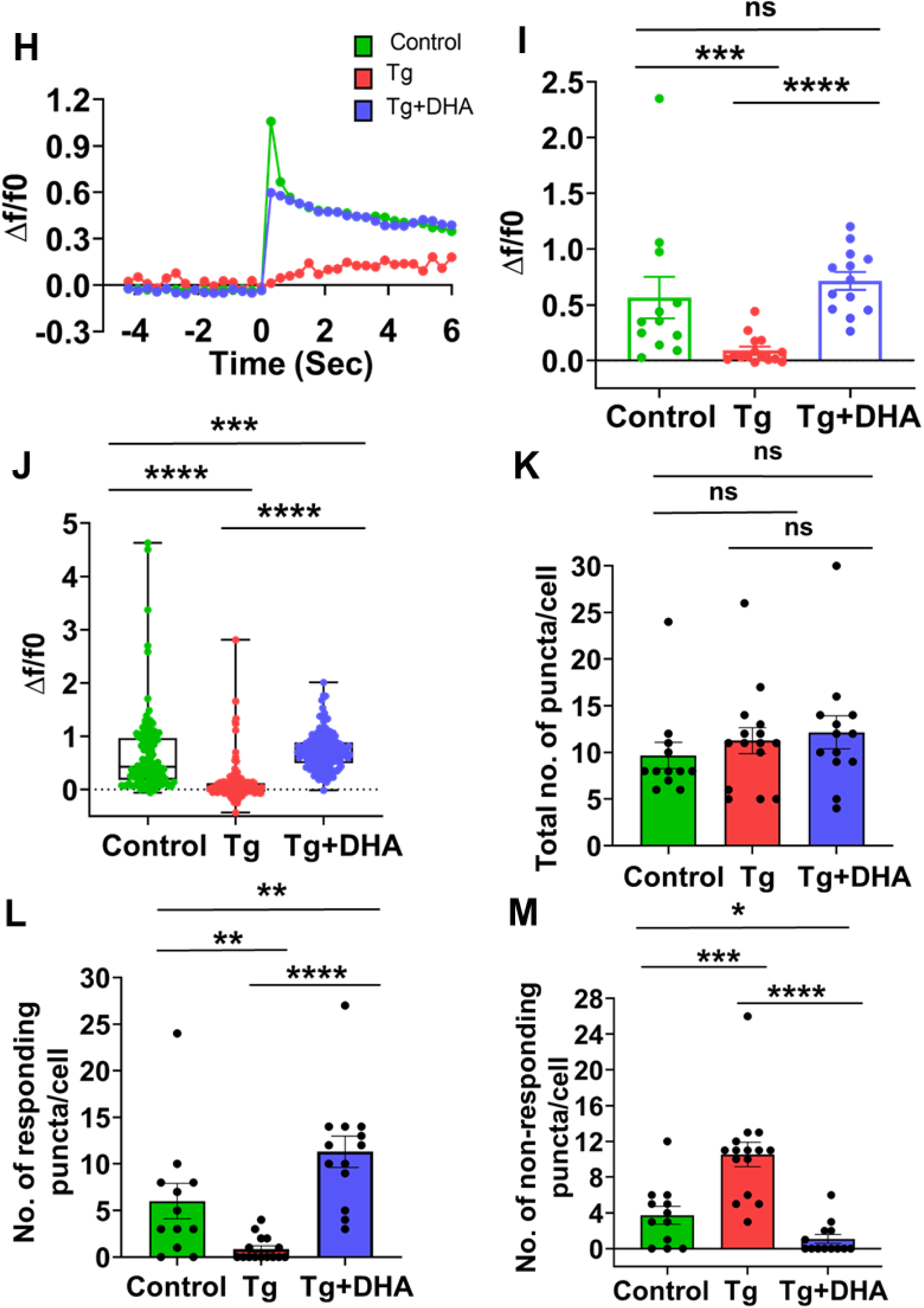
DHA rescues the phenotype by reducing ER stress in dense-core vesicles. **A** Treatment protocol: After plating, cells were serum-starved for 2 hours, followed by pre-incubating the cells with 10 µM DHA. After 24 hours, 10 nM Tg was given to the cells to induce ER stress in the DHA co-treated and Tg-alone cells. **B** Representative blots (left) and quantification (right) of protein levels of p-EIF2α/t-EIF2α. Beta tubulin served as a loading control (Student’s t-test, N=6). **C** Quantitative measure of NPY-mApple fluorescence intensities in Control, Tg treated, and DHA co-treated PC12 cells transfected with NPY-mApple showing basal and stimulated conditions, measured using a plate reader (Student’s t-test, N=5). **D** Quantification of total basal NPY-mApple fluorescence intensities (cell lysates and supernatants) (Mann-Whitney test, N=5). **E** Coomassie staining of constitutive secretion. **F** Immunoblot image of constitutive secretion marker Hsp-90. **G** Representative NPY-pHTomato (red) images in PC12 cells showing basal and KCl stimulated conditions (white arrows). **H** Representative time traces of mean NPY-pHTomato fluorescence intensities showing an average of all puncta per cell (green, control; red, Tg treated cells; blue, DHA co-treated cells). **I** Quantitative analysis of the maximum Δf/f0 in control, Tg, and DHA co-treated PC12 cells (Mann-Whitney test, Control, n=12; Tg, n=15; Tg+DHA, n=13; N=3). **J** Quantitative analysis showing the maximum Δf/f0 of all NPY-pHTomato puncta from all cells (Mann-Whitney test, Control, x=117; Tg, x=171; Tg+DHA, x=161; N=3). Data are shown as mean ± SEM. **(K-M)** Quantification of the total number of NPY-pHTomato puncta/cell (Mann-Whitney test) **(K)**, No. of responding NPY-pHTomato puncta/cell (Mann-Whitney test), **(L)** No. of non-responding NPY-pHTomato puncta/cell (Mann-Whitney test) **(M)** considering a minimum of Δf/f0 of 0.3 upon KCl stimulation as a response, (Control, n=12; Tg, n=15; Tg+DHA, n=13; N=3). (‘N’ denotes the number of biological replicates and ‘, n’ is the number of cells taken for quantification, ‘x’ is the number of puncta analyzed) Data are shown as mean ± SEM. (*P < 0.05; **P < 0.01; ***P<0.001; n.s. statistically non-significant).

**Figure 5:**
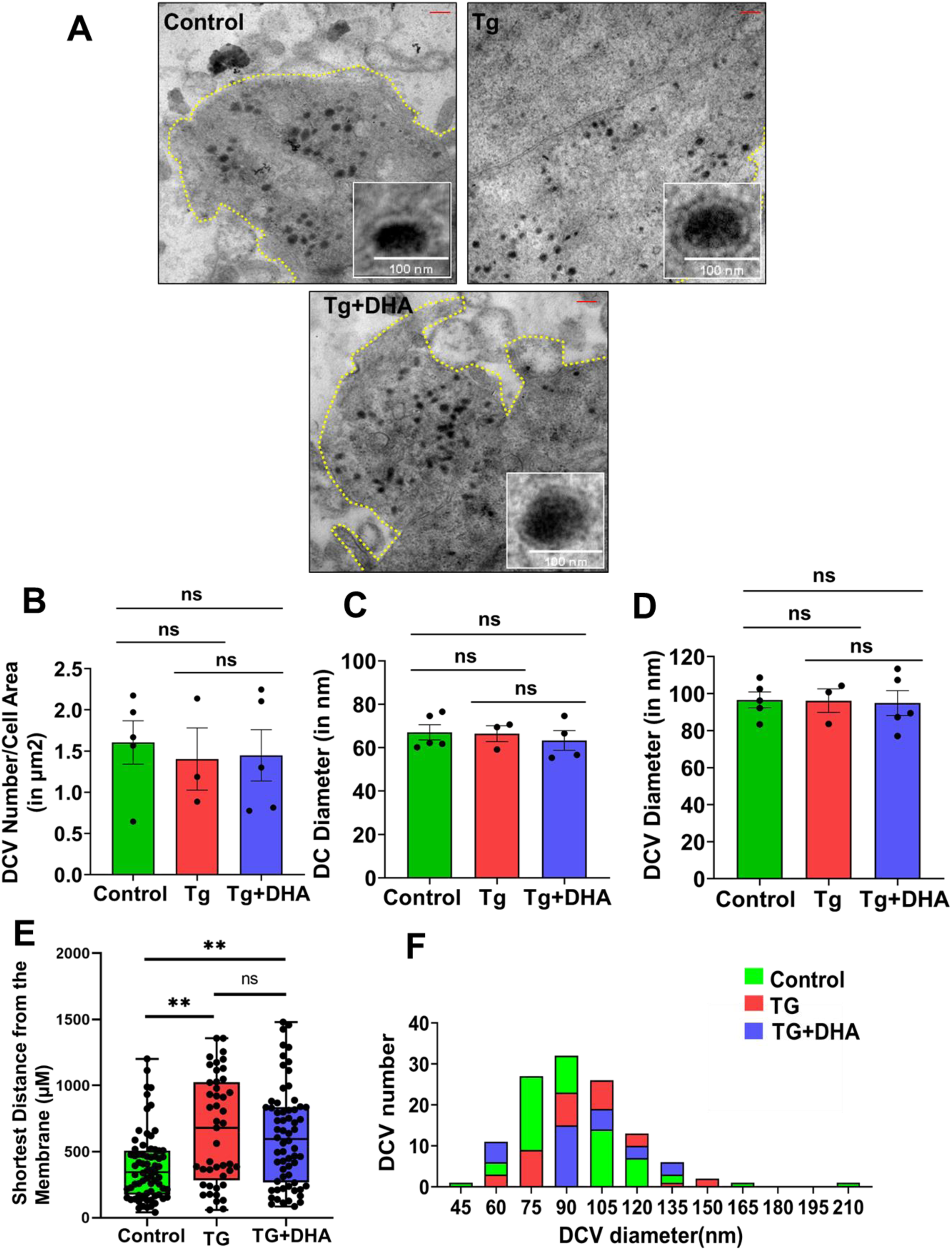
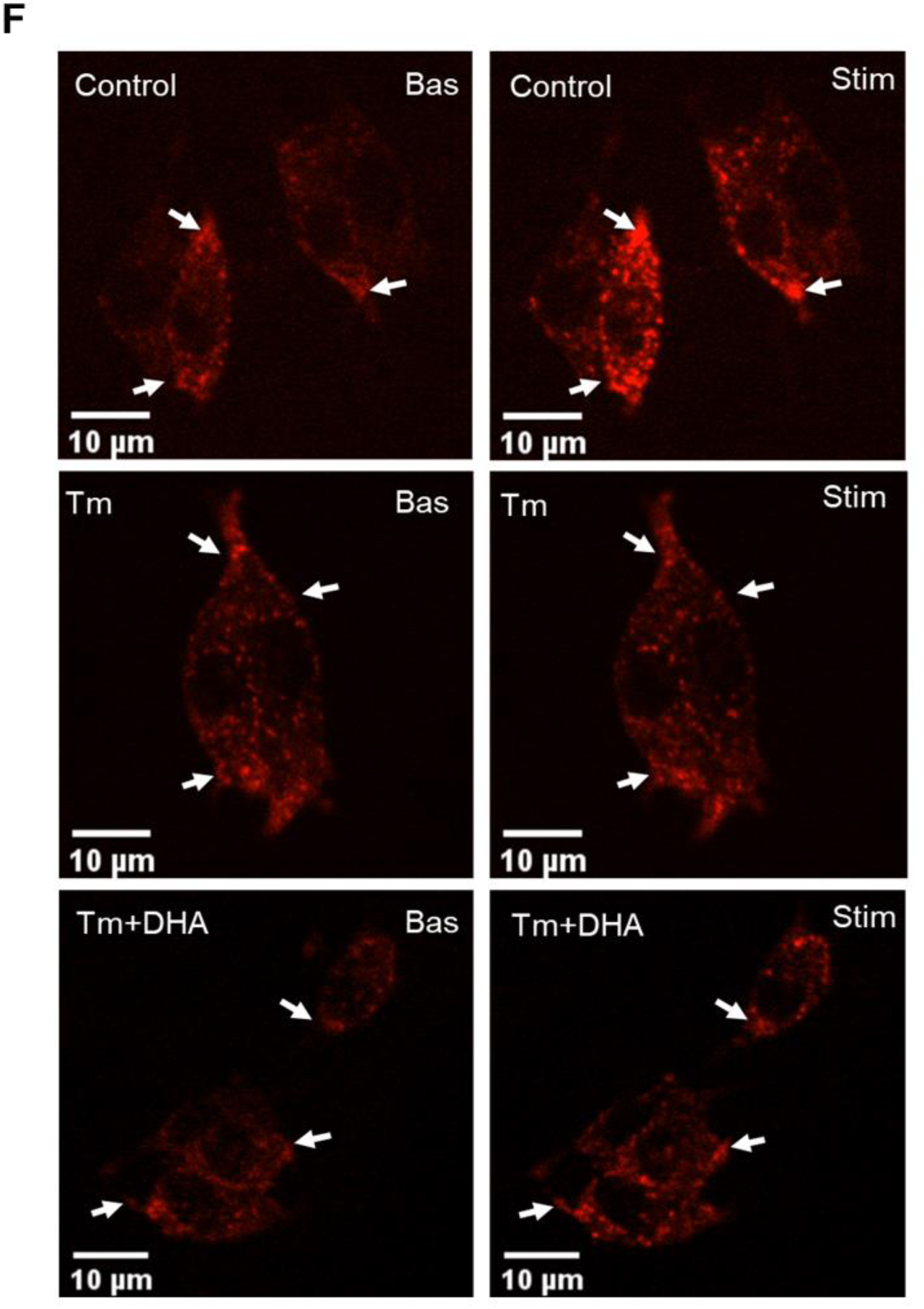
Transmission electron microscopy of PC12 cells. **A** Representative Transmission electron micrograph showing electron-dense vesicles. **B** Quantification of The number of DCVs/µM^2^ of the cell surface area (Student’s t-test; Control, n=5, Tg, n=3, Tg+DHA, n=5; N=2) **C** Quantification of dense core diameter (Student’s t-test; Control, n=5, Tg, n=3, Tg+DHA, n=5, N=2). **D** Quantification of dense core vesicle diameter (Student’s t-test; Control, n=5, Tg, n=3, Tg+DHA, n=5, N=2). Shows no significant difference. Quantification of DCV distance from the nearest plasma membrane, showing significant difference in DCV localization from the membrane. (Mann Whitney; n=5, Tg, n=3, Tg+DHA, n=5; N=2) **F.** Histogram of DCV Diameter Distribution (‘N’ denotes the number of biological replicates, and ‘n’ is the number of cells taken for quantification) Data are shown as mean ± SEM. (*P < 0.05; **P < 0.01; ***P<0.001; n.s. statistically non-significant).

We studied the pathway connected to ER stress inhibition and DCV exocytosis and biogenesis using DHA. Interestingly, DHA restored the impaired regulated exocytosis of DCV in Tg-treated cells to near control levels, as seen from the fluorescent-based assay (Fig. 4C, 4D). We later checked for the signature abundance of the constitutively secreted proteome by Coomassie staining the gel (Fig. 4E) and also with a specific constitutive secretory marker HSP-90 by immunoblotting at basal conditions without stimulation (Fig. 4F) (Guo et al., 2017) (Hummer et al., 2017). This ruled out impediments in the constitutive secretion phenomena.

Furthermore, we also observed an overall shift in the dynamics of DCV exocytosis as that of control cells upon DHA co-treatment (Fig. 4G, 4H), (Supp video 3) We also observed that the relative responsiveness of the cells to stimulus-coupled secretion was restored, as evident from the increased quanta of responsive NPY-pHTomato puncta upon DHA co-treatment (Fig. 4)(I-M). Hence, DHA treatment as an attenuator of ER stress does cause a change in DCV dynamics towards normal conditions along with the reversal of the impaired regulated secretion phenotype as seen in the Tg-treated cells. These experiments convey the specificity of the ER stress-associated perturbed regulated exocytosis, besides flagging the therapeutic potential of DHA where ER stress causes compromised regulated exocytosis.

### Tg treatment does not change the Size and Number but alters the spatial localisation of DCV

In light of severe disruption in DCV exocytosis, we wanted to estimate the size and distribution of the vesicles. TEM experiments followed by morphometry suggested no significant differences in the DCV size and number (Fig A-D). However, DCVs were significantly closer to the plasma membrane in the control cells than the ER-stressed counterpart (Fig E).

### Tunicamycin impedes DCV exocytosis, similar to Thapsigargin

To rule out that the observed cellular effect is a consequence of Tg, not ER stress, we tested another ER stressor with a different mechanism of action, Tunicamycin (Tm) (Guha et al., 2017). Tm also decreases cell viability without driving the cell towards apoptosis at 0.05 µg/ml dose (Fig. 6) (A, B), so for further experiments, this dose was chosen as the treatment dose. Tm upregulates the activity of Eif2α and, similar to Tg, impedes the secretion of DCV, which is subsequently restored to normal level upon DHA treatment. (Fig. 6) (A-L) (Supp. Video 6)

**Figure 6:**
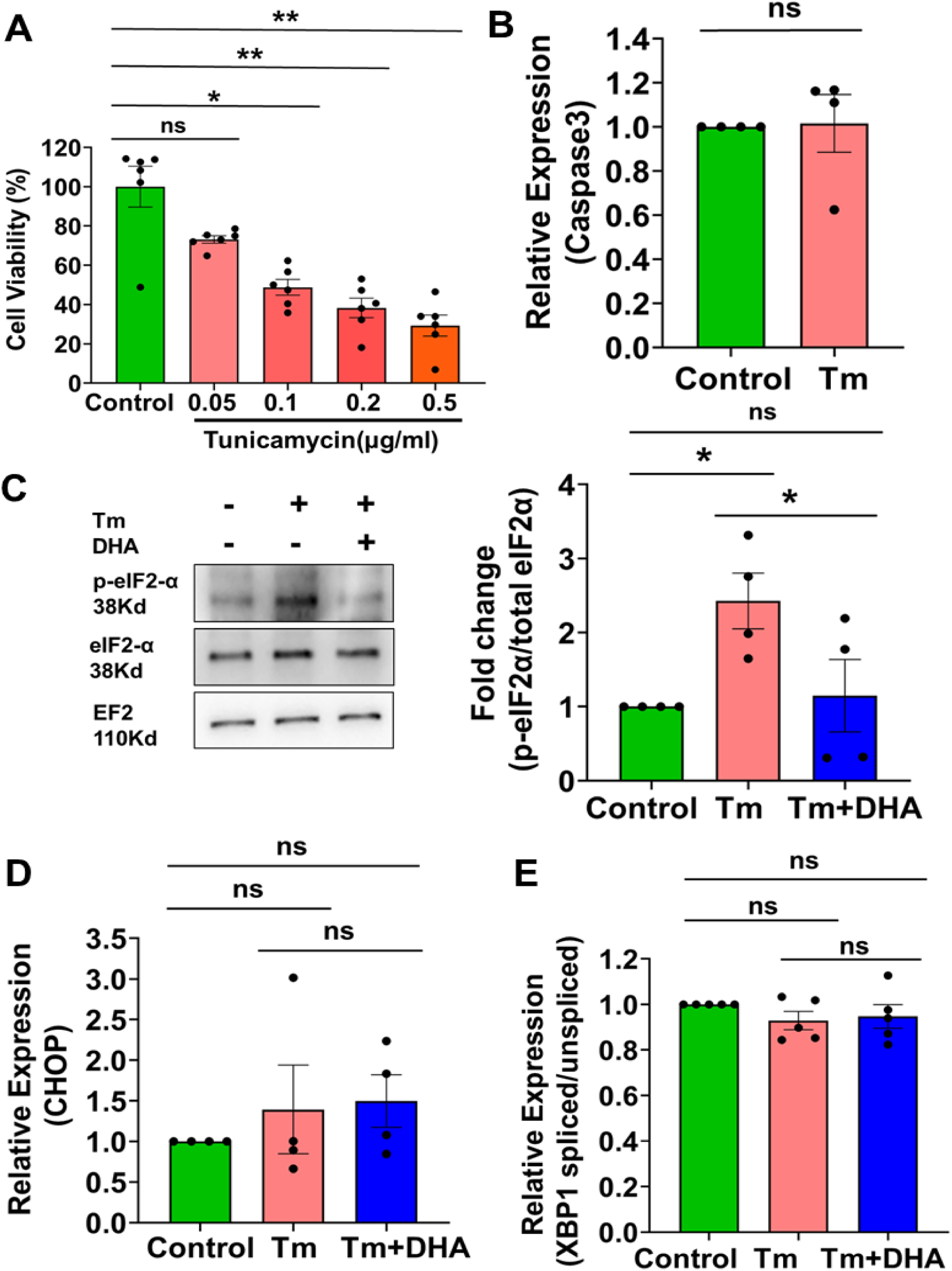
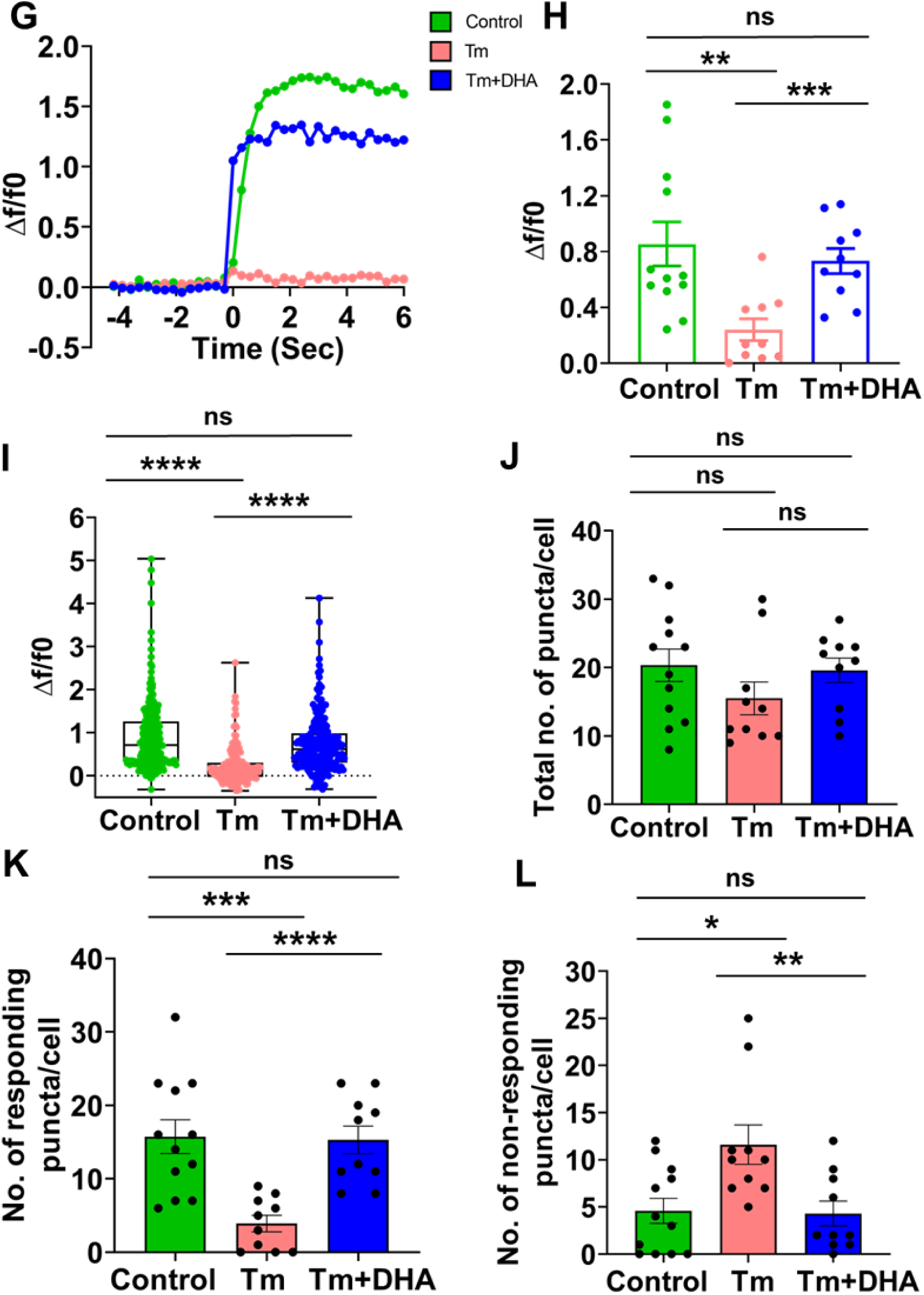
DCV exocytosis is impaired in tunicamycin-induced ER stress in PC12 cells. **A** Cell viability assay for different Tm doses showed greater than 70% cell viability in 0.05 µg/ml dosage (ANOVA, N=6). **B** Transcript levels of Caspase 3 in Control and Tm treated cells (Mann-Whitney test, N=4). **C** Representative immunoblot showing p-eIF2α and t-eIF2α (left) expression and its corresponding Quantification (right) in Control and cells treated with Tm (Student’s t-test, N=4). EF2 served as a loading control. **D** Quantitative analysis showing Transcript levels of CHOP (Mann-Whitney test, N=4). **E** Quantitative analysis showing Transcript levels of Xbp1 spliced/unspliced (Student’s t-test, N=5). **F** Representative NPY-pHTomato (red) image in PC12 cells showing basal and KCl stimulated conditions (white arrows). **G** Representative time traces of mean NPY-pHTomato fluorescence intensities showing an average of all puncta per cell (green, control; red, Tm treated cells; blue, DHA co-treated cells). **H** Quantitative analysis of the maximum Δf/f0 in control, Tm, and DHA co-treated PC12 cells (Mann-Whitney test, Control, n=12; Tm, n=10; Tm+DHA, n=10; N=3). **I** Quantitative analysis showing the maximum Δf/f0 of all NPY-pHTomato puncta from all cells (Mann-Whitney test, Control, x=244; Tg, x=155; Tg+DHA, x=196; N=3). **J** Quantification of the total number of NPY-pHTomato puncta/cell (Mann-Whitney test, Control, n=12; Tm, n=10; Tm+DHA, n=10; N=3) **K** Quantification of no. of responding NPY-pHTomato puncta/cell (Student’s t-test, Control, n=12; Tm, n=10; Tm+DHA, n=10; N=3). **L** Quantification of no. of non-responding NPY-pHTomato puncta/cell (Mann-Whitney test, Control, n=12; Tm, n=10; Tm+DHA, n=10; N=3) considering a minimum of Δf/f0 of 0.3 upon KCl stimulation as a response. (‘N’ denotes the number of biological replicates, and ‘n’ is the number of cells taken for quantification, x= no of puncta analyzed) Data are shown as mean ± SEM. (*P < 0.05; **P < 0.01; ***P<0.001; n.s. statistically non-significant).

### DHA reverses impediment to regulated secretion in SLVs

In line with our previous observation, we checked for the effect of DHA co-treatment in the exocytosis of SLVs in response to 100 mM KCl in ER stressed cells. Although there was a subtle restoration of impaired regulated secretion of SLVs in DHA co-treated ER-stressed cells, the magnitude of restoration was not in the order of DCV exocytosis (Fig. 7)(A-C), Supp video 4). We also observed that most of the Synaptophysin-pHTomato puncta showed non-responsive behaviour in Tg-treated cells, which were rescued to near control levels under co-treatment with DHA (Fig. 7F, 7G). We report that DHA treatment as an inhibitor of ER stress also causes a change in SLV dynamics towards normal or wild-type conditions along with the reversal of the impaired regulated secretion phenotype as seen in the ER-stressed cells.

**Figure 7:**
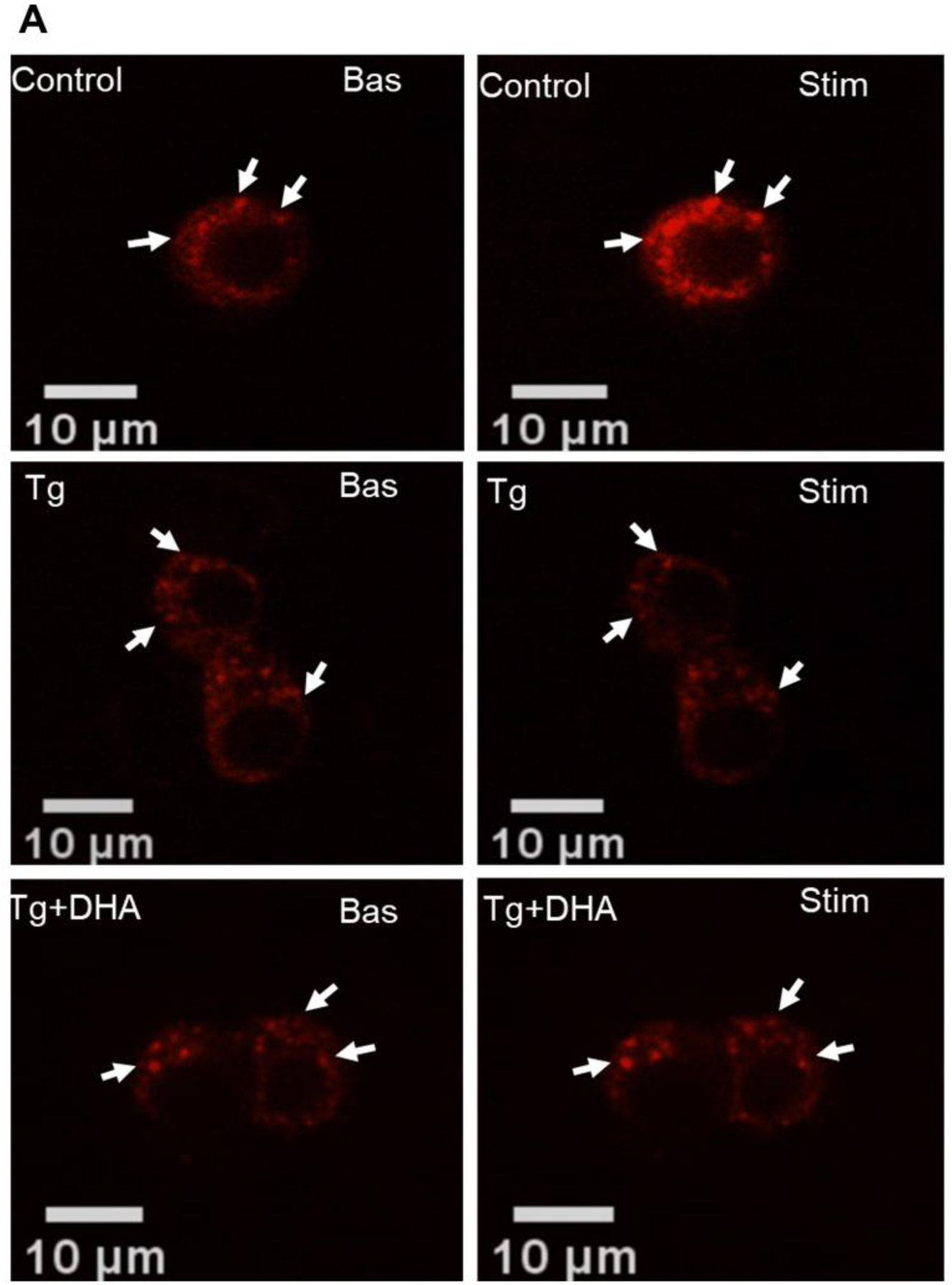
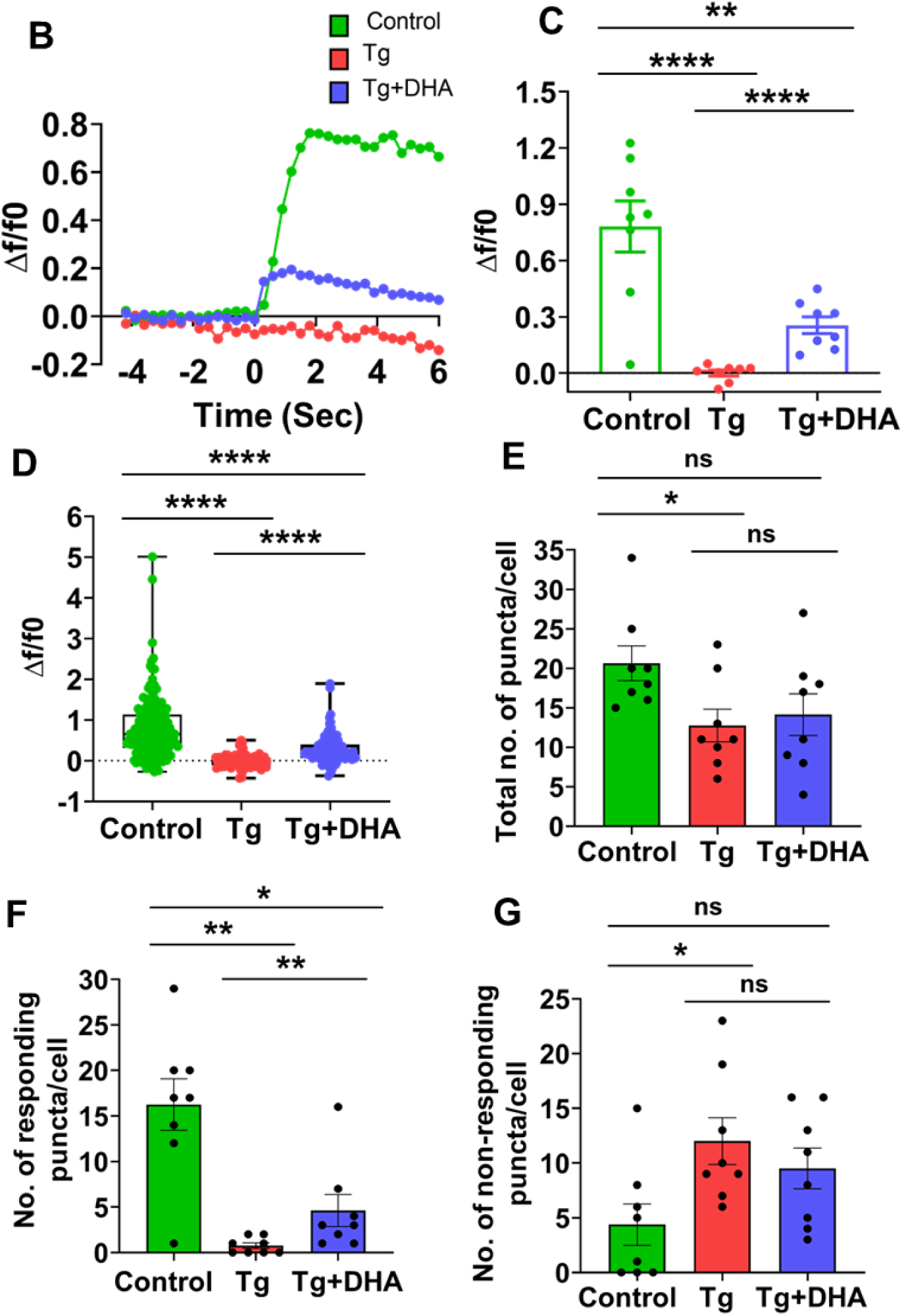
DHA rescues the phenotype by reducing ER stress in synaptic vesicles. **A** Representative image of Synaptophysin-pHTomato (red) in PC12 cells showing basal and KCl stimulated conditions (white arrows). **B** Representative time traces of mean Synaptophysin-pHTomato fluorescence intensities showing an average of all puncta per cell (green, control; red, Tg treated cells; blue, DHA co-treated cells). **C** Quantitative analysis of the maximum Δf/f0 in Control, Tg, and DHA co-treated PC12 cells (Student’s t-test, Control, Tg, Tg+DHA, n=8, N=3). **D** Quantitative analysis showing the maximum Δf/f0 of all Synaptophysin-pHTomato puncta from all cells (Mann-Whitney test, Control, x=165; Tg, x=96; Tg+DHA, x=113; N=3). Data are shown as mean ± SEM. **(E-G)** Quantification of total number of Synaptophysin-pHTomato puncta/cell (Student’s t-test) **(E)**, No. of responding Synaptophysin-pHTomato puncta/cell (Mann-Whitney test) **(F)** No. of non-responding Synaptophysin-pHTomato puncta/cell (Student’s t-test) **(G)** considering a minimum of Δf/f0 of 0.3 upon KCl stimulation as a response, (Control, Tg, Tg+DHA, n=8; N=3). (‘N’ denotes the number of biological replicates, and ‘n’ is the number of cells taken for quantification, x= no of puncta analyzed) Data are shown as mean ± SEM. (*P < 0.05; **P < 0.01; ***P<0.001; n.s. statistically non-significant).

### ER stress alters the abundance of signature genes associated with exocytosis

We conducted experiments to understand the molecular basis for impaired exocytosis by pharmacological ER stress triggered by Thapsigargin. Previous reports suggested blunted expression of key exocytotic genes such as *Snap25* in metabolic stress, consequently causing failure of exocytosis (Liew et al., 2010). Hence, we hypothesized that a diminished abundance of SNAP25 levels could cause impaired exocytosis in our ER stress cell model. Consistent with our hypothesis, SNAP25 levels were strikingly diminished in Tg-treated cells, which were restored to near-normal levels upon DHA co-treatment, as seen in (Fig. 8A, 8B). These results were consistent with other ER stressor Tm (Supp Fig.4). Surprisingly, these findings were contradicted by an increased CREB (cAMP Response Element Binding Protein) activity which was persistent upon DHA co-treatment (Fig. 8C). It is noteworthy that CREB is an established positive regulator of SNAP25 expression (Jin et al., 2021; Wang et al., 2019). This unexpected finding led us to test the expression profile of *Atf4*, a competitive inhibitor of CREB (also known as CREB 2) and a downstream effector molecule of the PERK-EIF2α axis (Adjibade et al., 2017; Dey et al., 2012; Liew et al., 2010; Smith et al., 2020). We observed an increase in the mRNA and protein level of ATF4 in the ER-stressed cells, which reverted towards control levels upon DHA co-treatment (Fig. 8D, 8E) (Supp. Fig 4). Our results indicate a possible negative modulatory role of ATF4 on SNAP25 expression by interfering with CREB activity. Furthermore, experiments on the specificity of altered abundances for other molecular players, such as key SNARES, tethers and rabs, were also tested without significant differences (Supp Fig. 3) (A-F).

**Figure 8:**
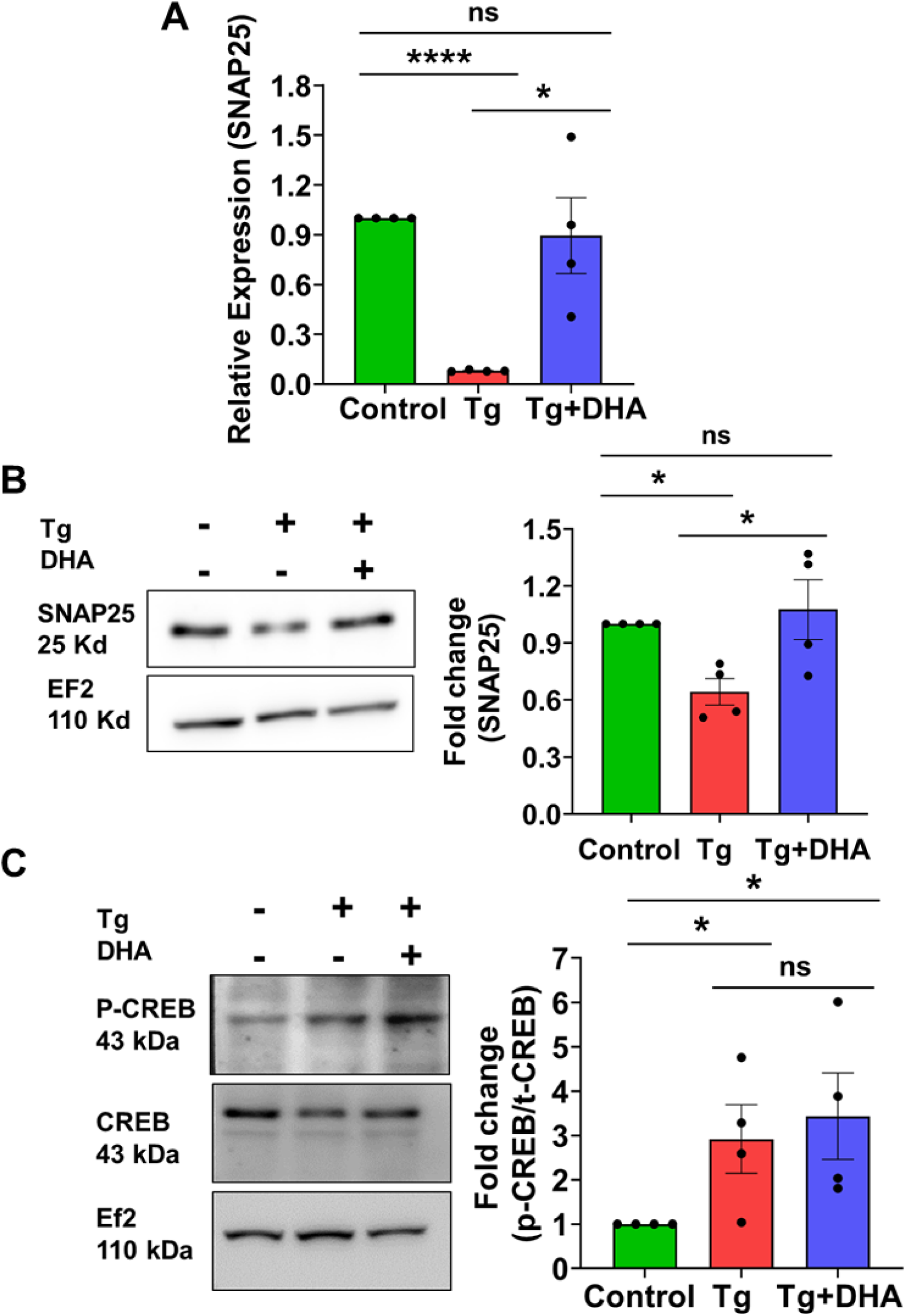
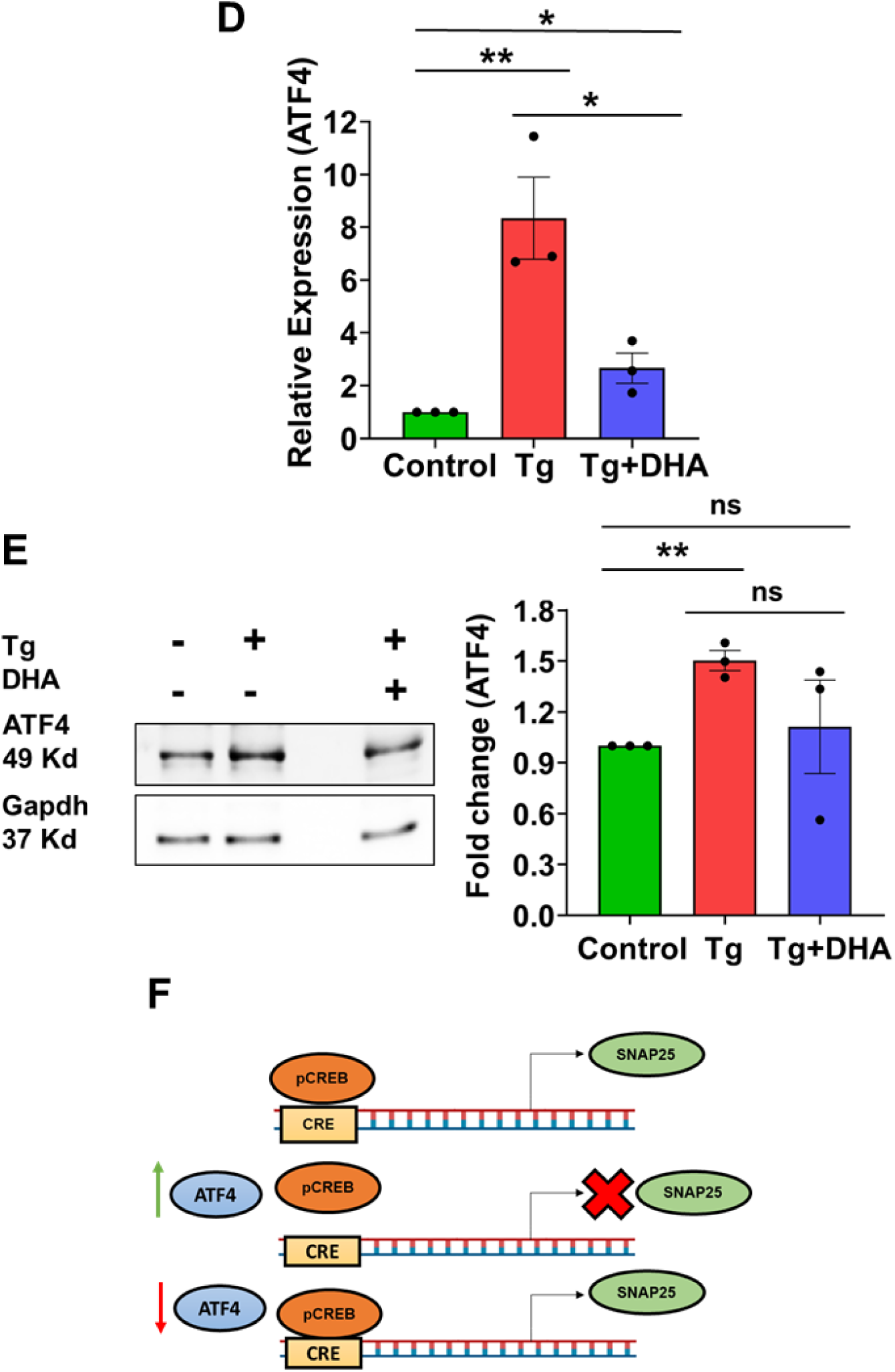
Change in the expression of exocytosis regulating gene SNAP25: **A** Quantification showing the transcript levels of *Snap25*, a t-SNARE which helps in DCV exocytosis by Q-RT PCR (Student’s t-test, N=4). **B** Representative blots of SNAP25 (left) and their corresponding quantification. EF-2 served as a loading control. (Student’s t-test, N=4). **C** Representative blots of p-CREB and t-CREB (left) and their corresponding quantification showing the ratio of p-CREB/t-CREB (right). EF-2 served as a loading control. (Student’s t-test, N=4). **D** Quantification showing *Atf4* transcript levels in Q-RT PCR (Student’s t-test, N=3). **E** Representative blots of ATF4 (left) and their corresponding quantification. GAPDH served as a loading control. (Student’s t-test, N=3) **F** Schematic representation of the transcriptional switch of CREB in the presence of ATF4. (‘N’ denotes the number of biological replicates) Data are shown as mean ± SEM. (*P < 0.05; **P < 0.01; ***P<0.001; n.s. statistically non-significant).

### ER, stress lowers the abundance of key granulogenic signatures

Given the well-documented evidence about the granulogenic potential of chromogranins, specifically Secretogranin II (SCGII) and Chromogranin A (CGA) (Kim et al., 2001; Lin et al., 2022), we wanted to test if there are any significant changes in the relative abundances of chromogranins in ER-stressed cells. Interestingly, CGA levels were significantly decreased at the mRNA and protein levels, which surprisingly remained unchanged in DHA co-treated cells (Fig. 9)(A-D). A similar trend is also observed at the transcript and protein levels for SCGII (Supp. Fig. 5)(A-B). These findings are interesting, given that CREB has a role in the transcriptional regulation of Chromogranins and other neuropeptides (Huttunen et al., 2002; Lonze and Ginty, 2002; Satoh et al., 2009). Concomitantly, there was enhanced colocalization of CGA with a lysosomal marker Lamp1 in lyso20-positive Tg-treated PC12 cells, which remained unchanged upon co-treatment with DHA (Fig. 9E). These findings were substantiated by the augmented levels of PDI (Protein Disulfide Isomerase), an ER resident chaperone known for its role in re-routing of misfolded proteins to the lysosomes (Fig. 9F) (Cha-Molstad et al., 2015; Wang et al., 2022; Wilkinson and Gilbert, 2004). In summary, these experiments suggested the depletion of major granins, known for their granulogenic functions, both transcriptionally and translationally, further accompanied by enhanced lysosomal localization, leading to a severe impediment in regulated DCV exocytosis in ER-stressed conditions.

**Figure 9:**
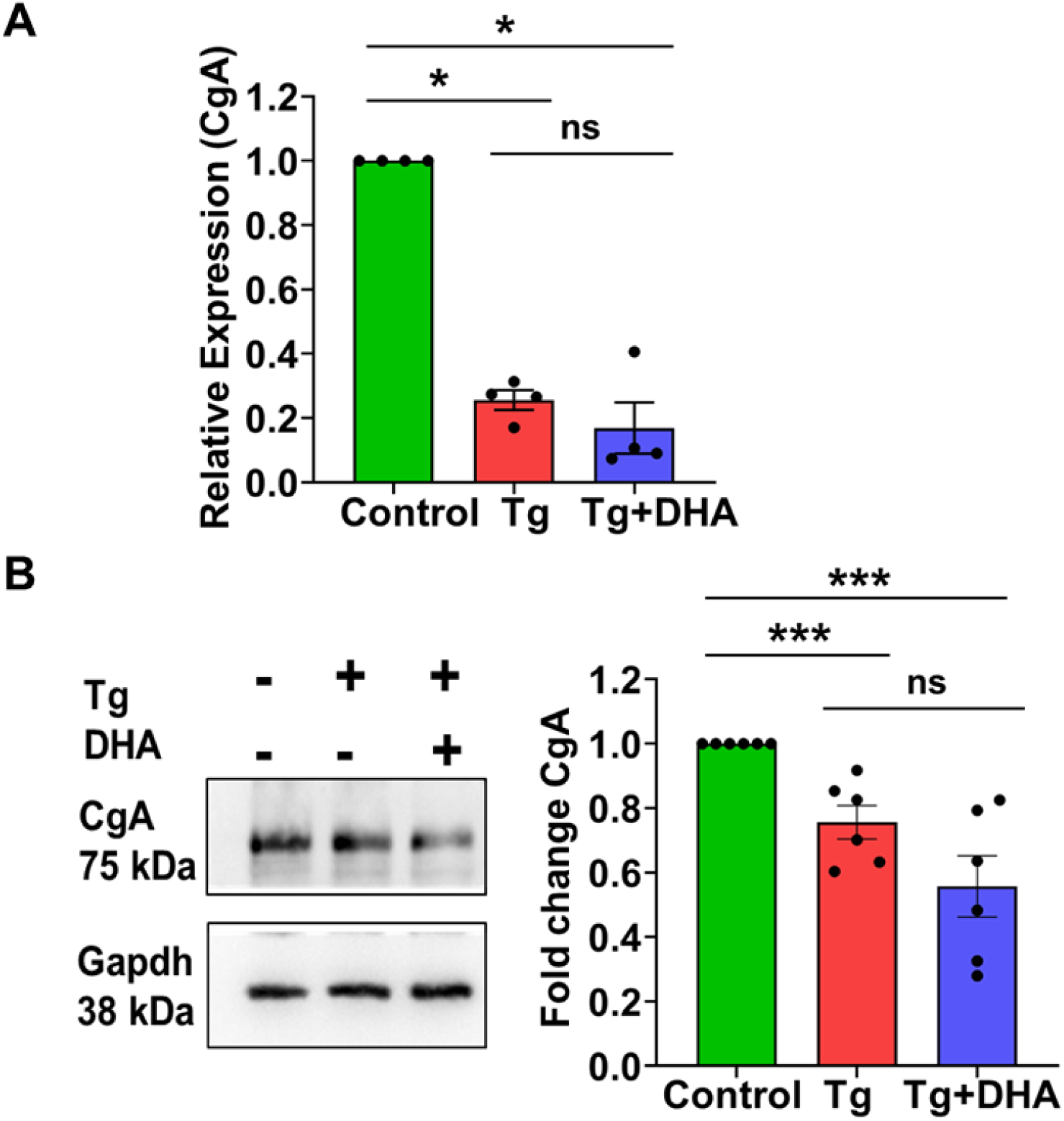
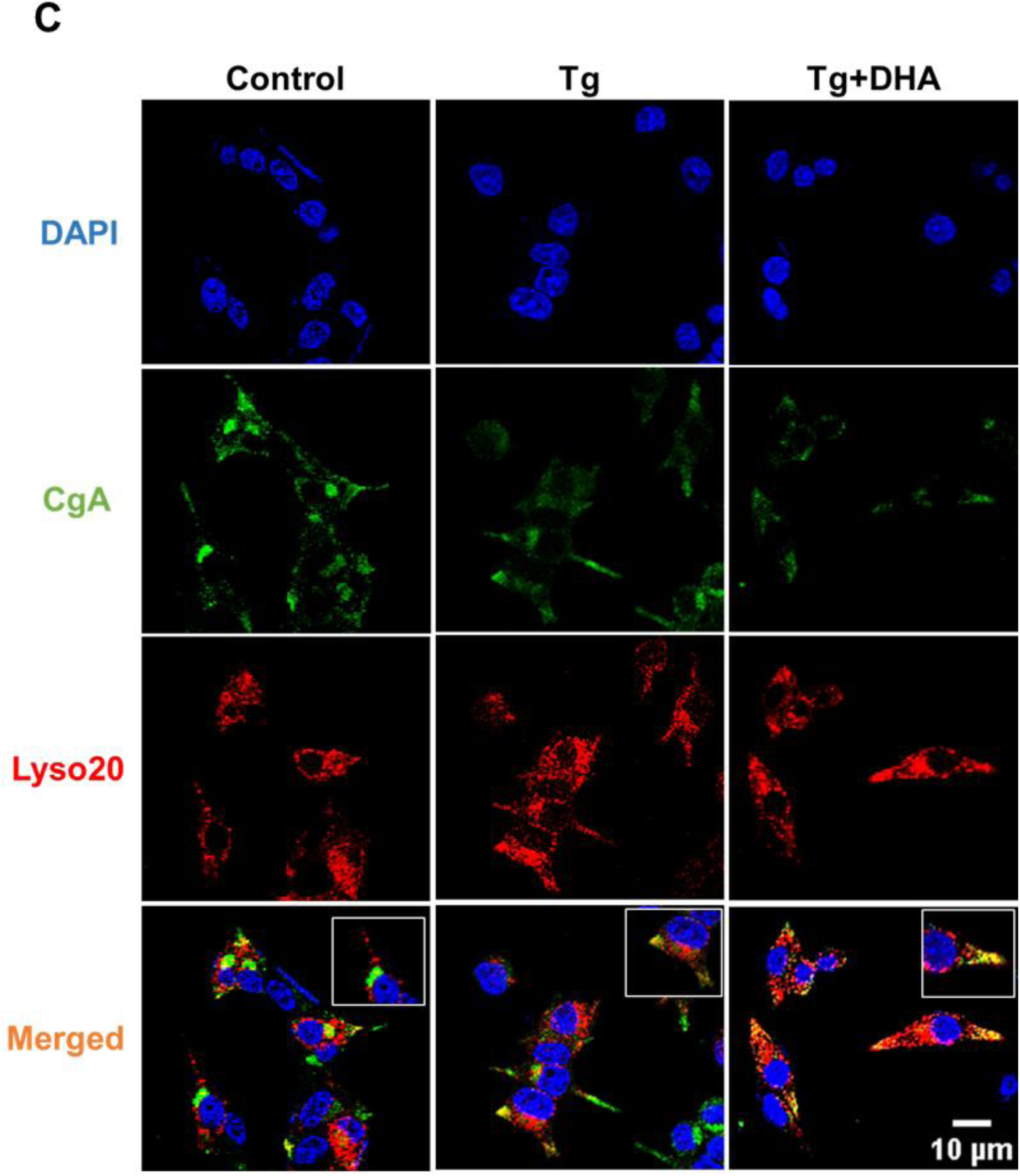
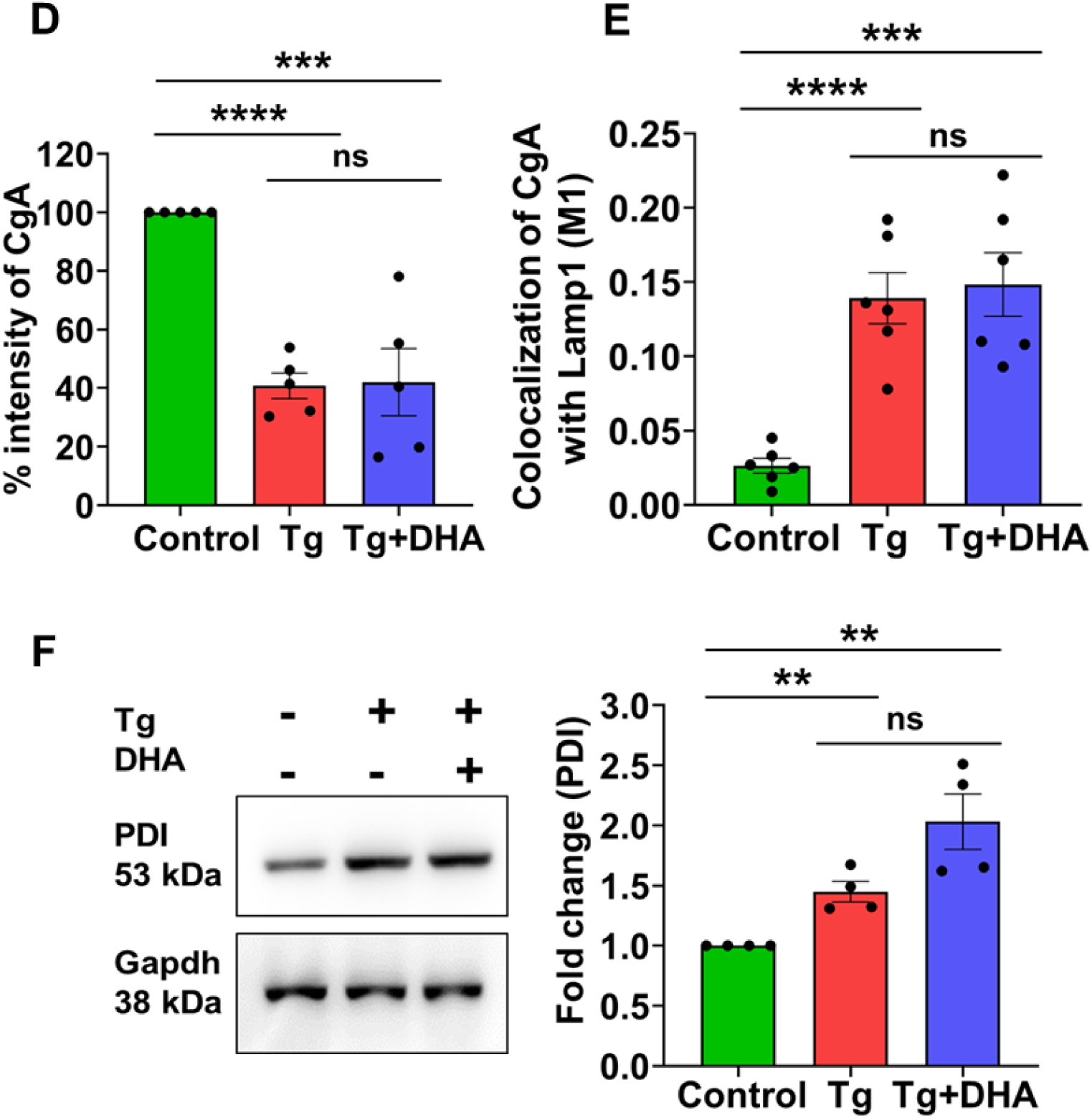
Change in DCV matrix proteins. **A** Transcript level of *Cga* in Control, Tg treated and DHA co-treated cells in Q-RT PCR, (Mann-Whitney test, N=4) **B** Representative blots of CGA (left) and Quantification of the protein level of CGA, (right). GAPDH served as a loading control. (Student’s t-test, N=6) **C** Representative immunofluorescence images showing DAPI (blue) and CGA (green) immunostaining in mCherry-Lyso20 (red) transfected PC12 cells, scale bar=10µm (2048*2048 pixels) **D** Quantification of CGA fluorescence intensity levels in Control, Tg treated and DHA co-treated cells (images quantified were captured in 512*512 pixels), (Student’s t-test, n=5; N=3) **E** Quantification showing Mander’s correlation coefficient between CGA and LAMP1 (Fraction of CGA overlapping LAMP1), (Student’s t-test, n=6, N=3). **F** Representative Blot of PDI (left) and Quantification of PDI expression (right) in western blot analysis. GAPDH served as a loading control. (Student’s t-test, N=4). (‘N’ denotes the number of biological replicates, and ‘n’ is the number of coverslips taken for quantification) Data are shown as mean ± SEM. (*P < 0.05; **P < 0.01; ***P<0.001; n.s. statistically non-significant).

**Figure 10:**
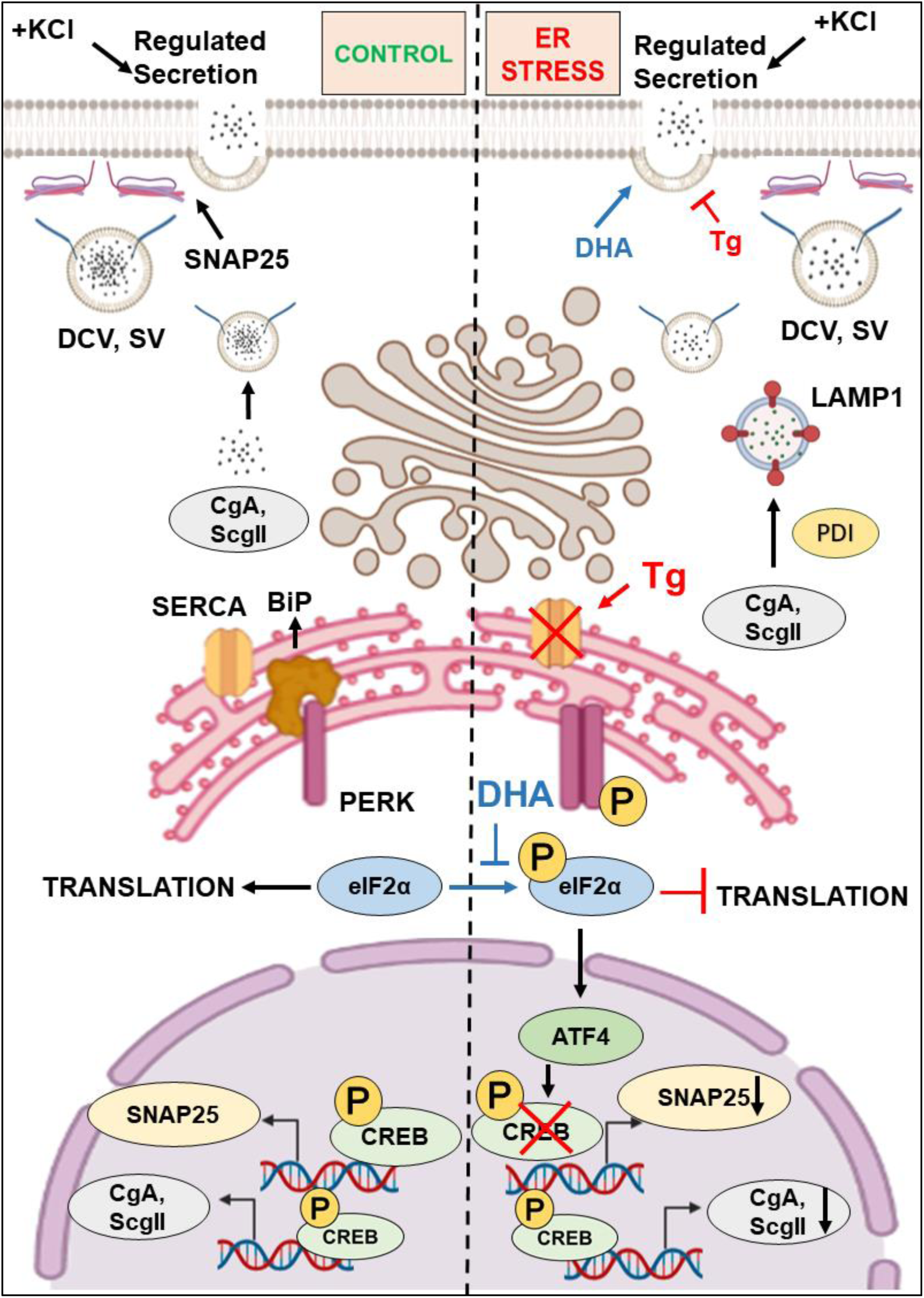
Plausible mechanism of action of impaired regulated secretion in ER-stressed cells. Thapsigargin-induced ER stress activates the PERK/EIF2α arm causing reduced expression of key exocytotic (SNAP25) and granulogenic (chromogranins) signatures via the CREB/ATF4 axis cumulatively responsible for the failure of regulated exocytosis.

## DISCUSSION

Previous studies have shown that a disruption in the release of components of dense core vesicles (DCVs) (Barranco et al., 2021) and synaptic-like vesicles (SLVs) (Zou et al., 2021) is tightly related to almost all neurodegenerative diseases. Regulated exocytosis displayed by DCVs and SLVs is a highly orchestrated major cell physiological pathway. DCVs and SLVs contain neuropeptides, monoamines and neurotransmitters essential for cell survival and growth (Birinci et al., 2020; Salio et al., 2006; Speese et al., 2007; Zhang et al., 2011), and the compromised release of its content creates a significant challenge to cellular physiology which ultimately leads to apoptosis, a consequence of which is neurodegeneration (Ulrich et al., 2002). The basic pathways related to the biogenesis of these vesicles are more or less understood. However, specific areas, such as crosstalk with other organelles and their behaviour during certain physiologically challenging conditions, such as metabolic stress, ER stress, or autophagy, are unexplored research avenues. We have pharmacologically modelled ER stress in professional neuroendocrine cells, such as PC12 and INS-1, without causing substantial cell death; hence, we call this ER stress, which was used as an ER stress paradigm for all subsequent experiments.

Interestingly, ER stress caused severe impairment in DCV exocytosis in two separate secretory cell lines, PC12 and INS-1 cells. Similar defects were also observed in SLV exocytosis, while no defects in the constitutive secretory pathway. To check the specificity of the ER stress pathway for its contribution to the impediment of regulated secretion, we rescued the cells from ER stress by using a well-known ER stress attenuator-an Omega 3 fatty acid, DHA-which also happens to be a neuroprotective compound (Begum et al., 2014, 2013, 2012b; Dyall, 2015). DHA co-treatment reversed eIF2α activation and strikingly negated the DCV exocytotic defects. Multiple ER stressors and attenuators in two independent cell lines substantiate our findings of impaired regulated secretion in an ER-stressed cell. Interestingly, stimulus-driven calcium extracellular influx via voltage-dependent ion channels, the major source for calcium-dependent DCV exocytosis, remains unchanged in Tg-treated cells compared to the control (Zorec, 1996) (Preissler et al., 2020) (data not shown).

To understand the molecular underpinning of ER stress-associated impaired exocytosis of DCV, we quantified exocytotic genes, which are fundamentally responsible for secretory vesicle exocytosis, expression switches of such regulatory genes are critical for active functional exocytosis. SNAP25 is one important membrane-associated t-SNARE required for the fusion of specialised secretory vesicles with the plasma membrane (Arora et al., 2017; Tsuboi and Fukuda, 2005; Walter et al., 2010). Such molecular switches are further regulated by master genes such as CREB (cAMP Response Element Binding protein), which is well-known for regulating SNAP25 expression (Jin et al., 2021; Liew et al., 2010; Wang et al., 2019). Intriguingly, impaired exocytosis in ER-stressed cells was correlated with reduced expression of SNAP25 levels, while several snares, tethers and rabs involved in exocytosis remain unchanged in our studies. We suspected CREB as an important candidate for such altered expression. However, to our surprise, these findings did not coincide with CREB activity, albeit substantiated by ATF4 levels, a downstream signalling molecule of one of the three UPR arms p-EIF2α and a well-known competitive inhibitor of CREB action.

Besides regulating the exocytotic switches, the ER-stressed cells also depicted reduced granins, specifically CGA and SCGII levels which remained lower upon DHA co-treatment. These findings are important given the well-established role of granins in Granulogenesis (granule formation) facilitated by assembly, condensation, and cargo aggregation, a hallmark of DCV maturation. These events are key as only mature DCVs are competent to undergo exocytosis. The chromogranin family mainly includes chromogranin A (CGA), chromogranin B (CGB), and secretogranin II (SCGII), which account for more than 90% of the DCV matrix proteins and are responsible for the accumulation and concentration of neuropeptides and calcium (Kim et al., 2001), (Mosley et al., 2007). Recent studies have also pointed out that biophysical events such as liquid-liquid phase separations of granins are also linked to granulogenic functions (Parchure et al., 2022). In parallel, expression switching of key granulogeneic factors can be potentially attributed to the augmented CREB activity, a well-known bidirectional transcriptional regulator of granins (Huttunen et al., 2002). A previous study has demonstrated that CREB downregulates CgA levels (Canaff et al., 1998) (Satoh et al., 2009). Furthermore, expression switching of granulogenic genes was also accompanied by the enhanced colocalization of CGA with a lysosomal marker. This finding can be potentially attributed to CGA being rerouted towards lysosomes by PDI, an ER-resident chaperone that routes some of the proteins to the autophagy-lysosomal pathway besides assisting in proper folding (Cha-Molstad et al., 2015; Wang et al., 2022).

Despite reduced expression of key granins, DHA treatment was able to reverse the altered regulated secretory phenotype. To understand this unexpected finding, we have performed transmission electron microscopy and morphometric analysis to quantify the vesicle number, dense-core size and DCV diameter, which were not significantly different. However, DCVS were more distantly located from the plasma membrane in Tg-stressed cells than control, an observation reverted with ER stress attenuator DHA (although not statistically significant). DCV localization away from the plasma membrane reduces the propensity of their release by decreasing the chances of vesicle docking and release.

We propose that these can be linked to the well-known and documented action of SNAP25, a key component of Stx1/SNAP25 acceptor complex (de Wit et al., 2009). Notably, SNAP25 levels were depleted in ER stressed cells, restored upon DHA treatment (Fig 8 A,B). However, Syt1 and Stx1 levels remain unchanged (Supp. Fig 3). Futuristically, studies can be envisaged to understand the role of ATF4 and its potential contributions as a master regulator in the transcriptional regulation of novel or known molecular switches associated with exocytosis and granule biogenesis.

In summary, for the first time, our study unleashes an impediment in the exocytosis of specialised secretory vesicles (DCVs & SLVs), although with different propensities in ER-stressed cells. To our knowledge, this is the first report on inter-organellar crosstalk amongst a stressed ER and specialized secretory vesicles. Compromised exocytosis of specialised secretory vesicles coupled with ER stress can act as a “double-edged sword “by causing enhanced detrimental effects due to cumulative disturbances of ER stress and the regulated secretory pathway. Furthermore, the reversal of DCV exocytosis by DHA, a well-known ER stress attenuator, can be of immense pharmacological significance, potentially paving the way to correct impaired DCV exocytosis in disease-associated ER-stressed conditions. These findings are of huge translational relevance given that a wide range of pathophysiological conditions such as neurodegeneration, diabetes, and cardiovascular diseases are associated with ER stress and neuromodulators/neurotransmitter released during regulated secretion with known neuroprotective functions such as combating ER stress itself (de Diego et al., 2020; Ozcan and Tabas, 2012). Although these are very interesting findings, we acknowledge certain limitations. For instance, the induced ER stress is artificial and unnatural, limited to cell culture experiments. In the future, It would also be interesting to look at the crosstalk of ER stress and investigate its association with the functional status of regulated exocytosis in animal models, under natural pathophysiological conditions such as diabetes, obesity, and neurodegenerative diseases such as HD/AD, given the known regulatory roles ER stress and DCV cargo in these diseases. (Cortès-Saladelafont et al., 2016; Thomas-Reetz and De Camilli, 1994).

ATF6: Activating Transcription Factor 6
ATF4: Activating Transcription Factor 4
BSA: Bovine Serum Albumin
CgA: Chromogranin A
CgB: Chromogranin B
CRE: cAMP-Response Element
CREB: cAMP-Response Element Binding Protein
DAPI: 4′,6-diamidino-2-phenylindole
DCV: Dense Core Vesicle
DHA: cis-4,7,10,13,16,19-Docosahexaenoic Acid
DMEM: Dulbecco’s Modified Eagle Medium
DMSO: Dimethyl Sulfoxide
DNA: Deoxyribonucleic Acid
EF2: Elongation Factor 2
eIF2α: Eukaryotic translation Initiation Factor 2
FBS: Fetal Bovine Serum
Gapdh: Glyceraldehyde-3-Phosphate Dehydrogenase
Grp78: Glucose-regulated protein 78
HEPES: 4-(2-hydroxyethyl)-1-piperazineethanesulfonic acid
HRP: Horse Radish Peroxidase
HS: Horse Serum
IRE1α: Inositol-requiring enzyme 1 alpha
LDCV: Large Dense Core Vesicle
MTT: 3-(4,5-dimethylthiazol-2-yl)-2,5-diphenyl tetrazolium bromide
NPY: Neuropeptide Y
PBS: Phosphate Buffered Saline
PC12: Pheochromocytoma 12
p-CREB: Phospho-cAMP-Response Element Binding Protein
PDI: Protein Disulfide Isomerase
PERK: Protein kinase R-like Endoplasmic Reticulum Kinase
PFA: Paraformaldehyde
p-IRE1α: Phospho-Inositol-Requiring Enzyme 1 alpha
PKA: Protein Kinase A
PLL: Poly-L-Lysine
PMSF: Phenylmethylsulfonyl fluoride
RIPA: Radioimmunoprecipitation Assay
RNA: Ribonucleic Acid
ScgII: Secretogranin II
SDS: Sodium Dodecyl Sulphate
SERCA: Sarcoendoplasmic Reticulum Calcium transport ATPase
SNAP25: Synaptosomal-Associated Protein
SNARE: SNAP Receptor
TBST: Tris Buffered Saline, with Tween-20
Tg: Thapsigargin
TGN: Trans-Golgi Network

## ACKNOWLEDGEMENTS

This study was supported by funds from the Raising Stars in Neuroscience award (formerly return home fellowship) by International Brain Research Organisation. A Ramalingaswami fellowship grant (BT/RLF/Re-entry/38/2016), an early career research grant from ICGEB, Trieste, and core funds from NBRC also funded the BSS lab towards this study. We thank NBRC core funds for providing salary support to this study’s authors. We would also like to thank Thomas Puccyadil and Dileep Vasudevan for their critical suggestions on the manuscript. BSS likes to thank his academic mentors, Profs. Nitish Mahapatra, Scottie Robinson, Alessandro Bartolomucci, and Sushil Mahata for their invaluable support. He would also like to thank Prof L.S. Shashidhara (IISER Pune/Ashoka University), Dr Meenakshi Munshi (former Advisor DBT) and Prof Anirban Basu, his career mentors. Lastly, BSS thanks all the colleagues at NBRC for helping to settle down amidst the pandemic and providing emotional and infrastructural support to our new lab.

## AUTHOR CONTRIBUTIONS

M.M, C.M, S.S, V.G, AJ and SD performed experiments. M.M. and BSS analysed the data and wrote the manuscript with inputs from all the co-authors. B.S.S conceptualized the study, designed the experiments with M.M, C.M, A.J and VG, and secured funding.

## Supplementary Information

**Figure S1:**
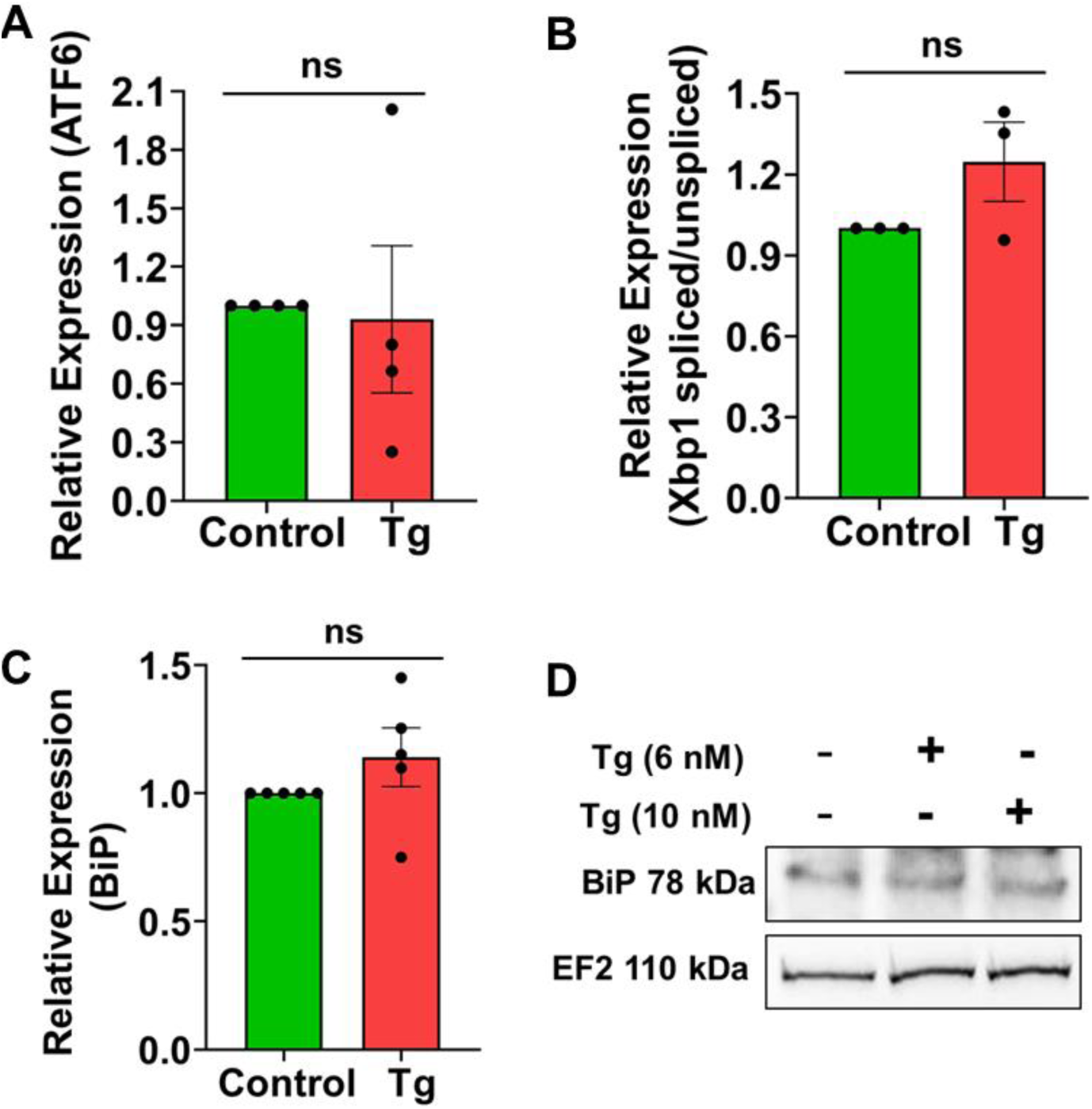
Thapsigargin causes ER Stress in PC12 cells. **A** Quantitative analysis showing Transcript levels of *Atf6* (Student’s t-test, N=4). **B** Quantitative analysis showing Transcript levels of *Xbp1* spliced/unspliced (Student’s t-test, N=3). **C** Quantitative analysis showing Transcript levels of *Bip* (Student’s t-test, N=4). **D** Immunoblot of BIP in Control and stressed cells. Data are shown as mean ± SEM. (‘N’ denotes the number of biological replicates) (*P < 0.05; **P < 0.01; ***P<0.001; n.s. statistically non-significant).

**Figure S2:**
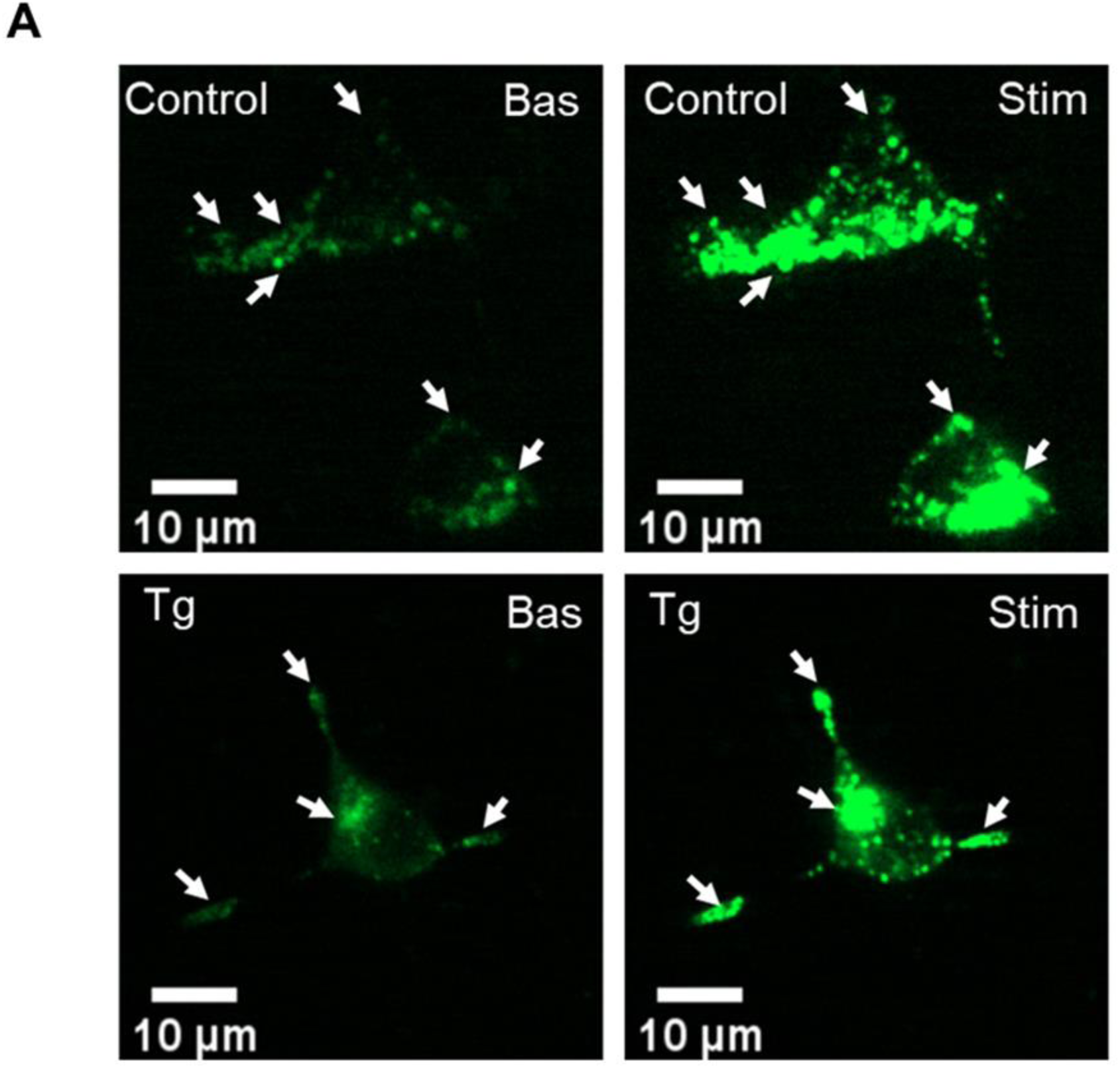
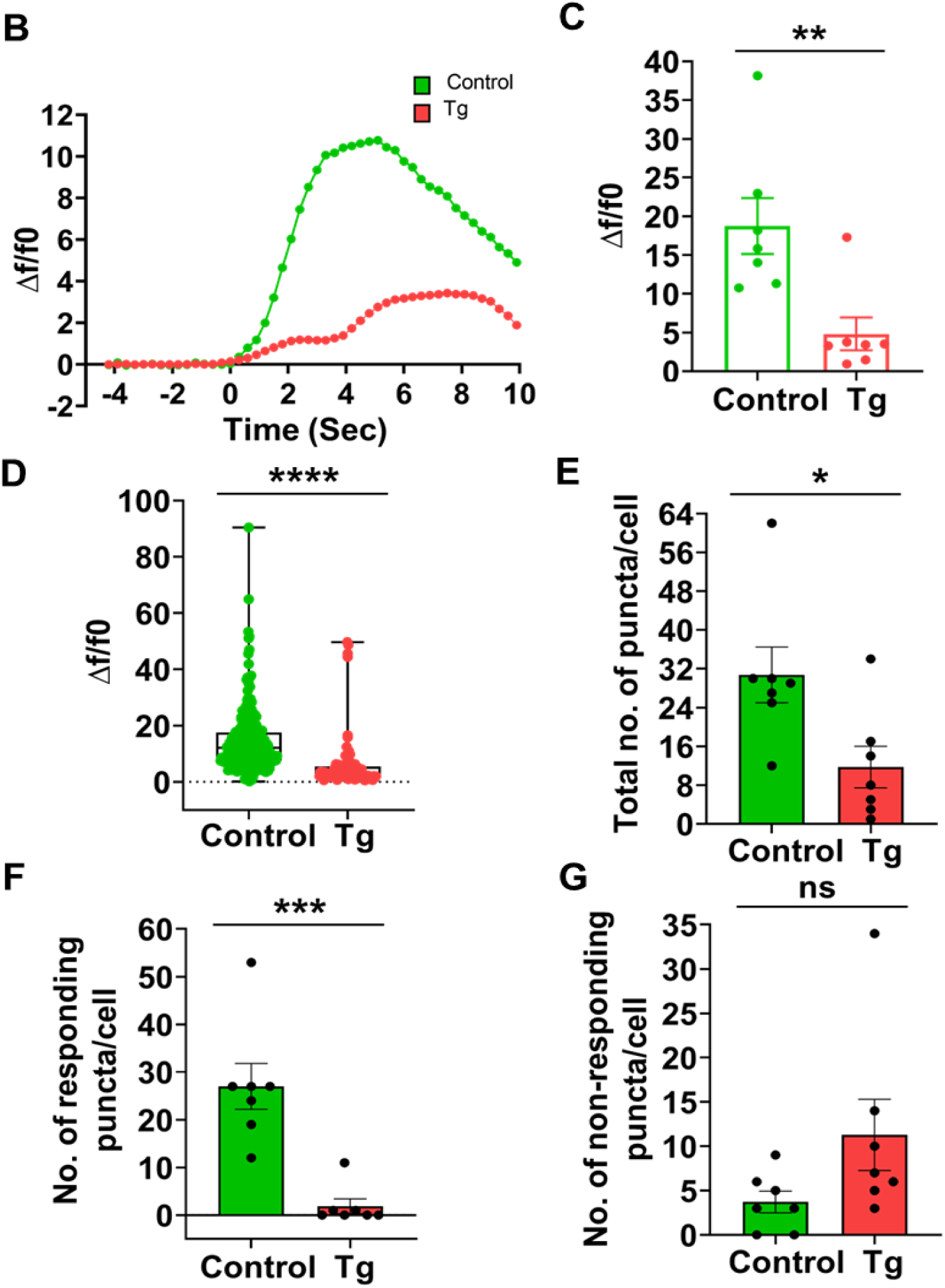
ER stress causes an impediment to regulated secretion in INS-1 cells. **A** Representative images of NPY-pHluorin (green) in INS-1 cells were captured from live cell imaging at 63X in a spinning disk at an interval of 300 ms (top, Control; bottom, Tg). **B** Representative time traces of mean NPY-pHluorin fluorescence intensities showing an average of all puncta per cell (green, control; red, Tg). **C** Quantitative analysis of the maximum Δf/f0 in Control and Tg-treated INS-1 cells (Mann-Whitney test, Control, Tg, n=7, N=3). **D** Quantitative analysis showing the distribution of Δf/f0 of all NPY-pHluorin puncta from all cells (Mann-Whitney test, Control, x=215; Tg, x=92). **(E-G)** Quantification of a total number of NPY-pHluorin puncta/cell (Mann-Whitney test) **(E)**, No. of responding NPY-pHluorin puncta/cell (Mann-Whitney test) **(F)** No. of non-responding NPY-pHluorin puncta/cell (Mann-Whitney test) **(G)** considering a minimum of Δf/f0 of 7 upon KCl stimulation as a response, (Control, Tg, n=7). (‘N’ denotes the number of biological replicates, and ‘n’ is the number of cells taken for quantification, x= no of puncta analyzed) Data are shown as mean ± SEM. (*P < 0.05; **P < 0.01; ***P<0.001; n.s. statistically non-significant).

**Figure S3:**
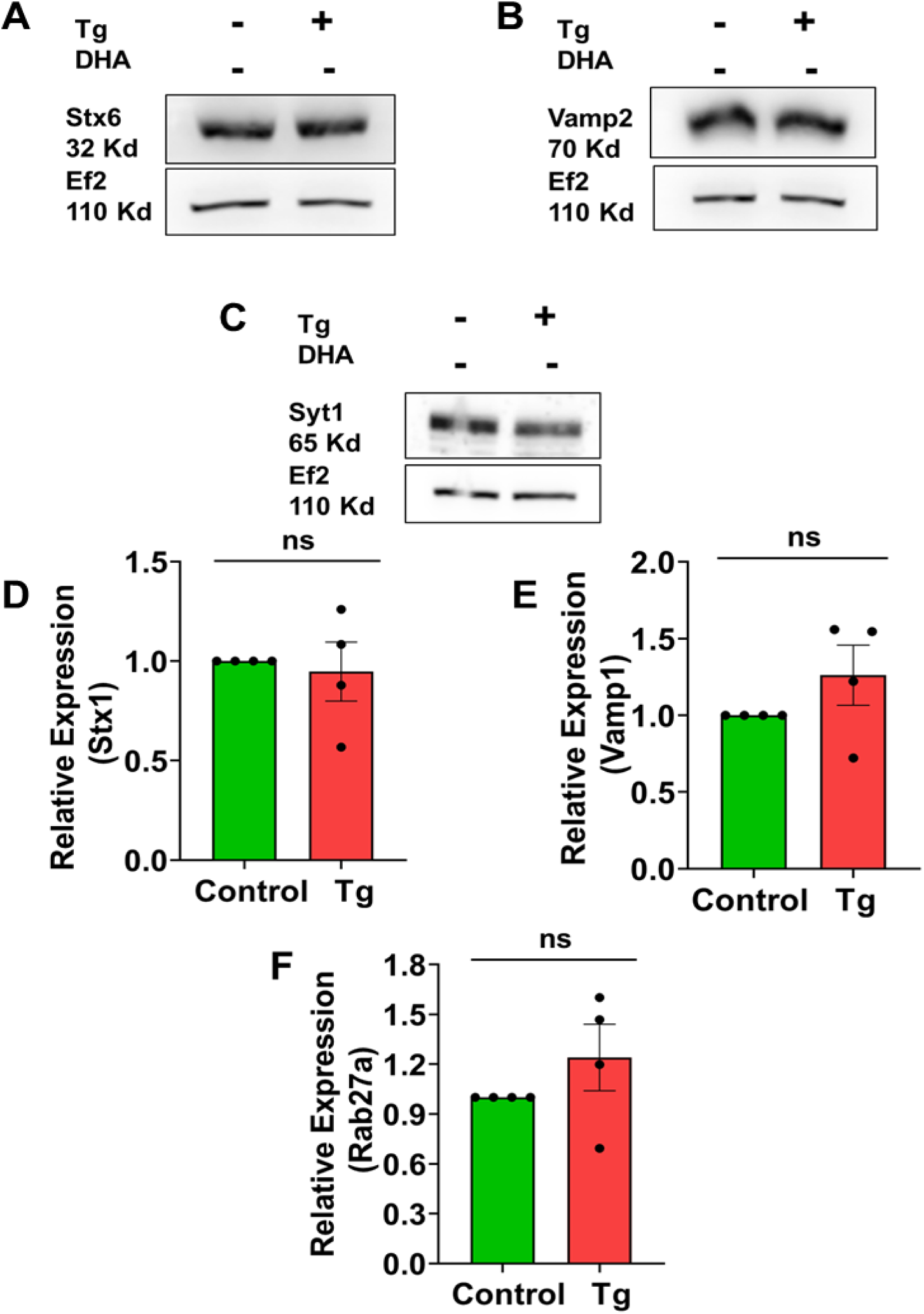
ER stress does not alter Exocytosis-related gene signature. **(A-C)** Representative immunoblots showing protein levels of STX6 **(A)**, VAMP2 **(B)** and SYT1 **(C).** EF2 served as a loading control. **(D-F)** Quantitative analysis showing Transcript levels of *Stx1* (Student’s t-test, N=4) **(D),** *Vamp1* (Student’s t-test, N=4) **(E)** and *Rab27a* **(F)** (Student’s t-test, N=4). (‘N’ denotes the number of biological replicates) Data are shown as mean ± SEM. (*P < 0.05; **P < 0.01; ***P<0.001; n.s. statistically non-significant).

**Figure S4:**
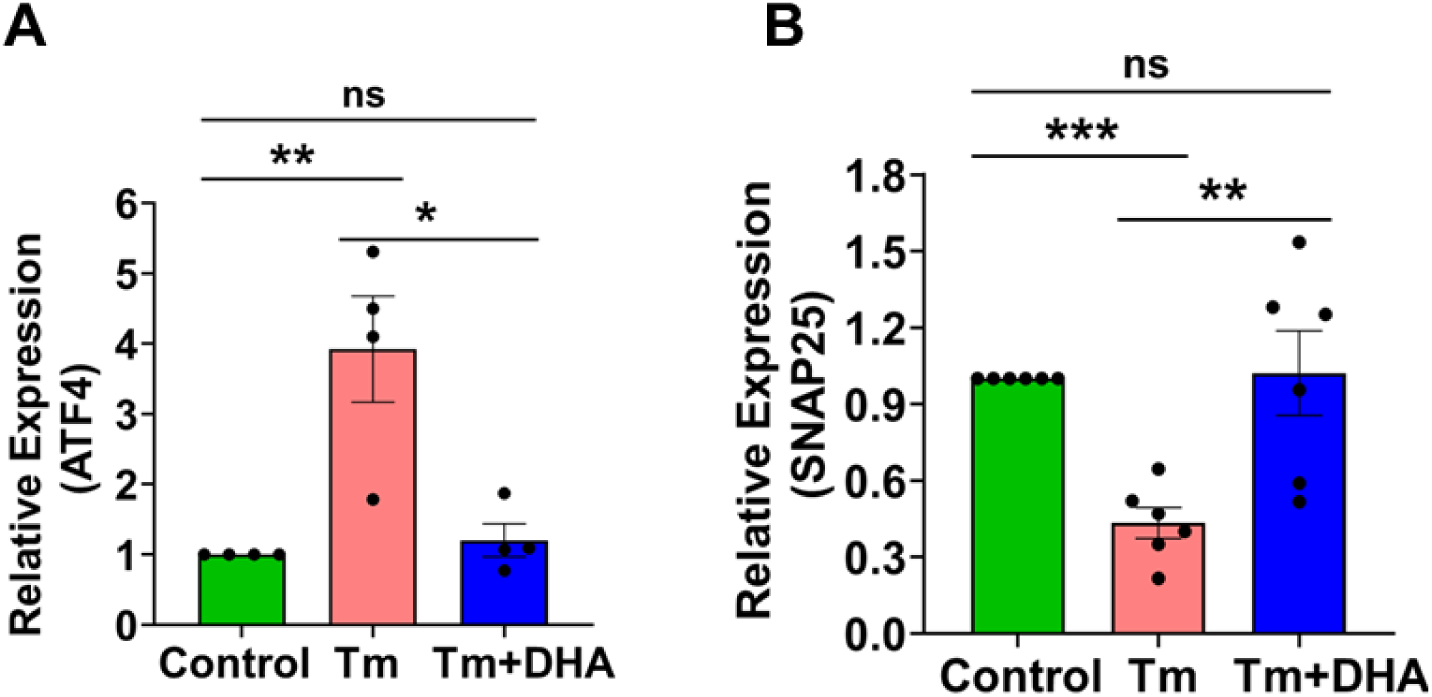
Tunicamycin treatment affects the expression of SNARE. **A** Quantitative analysis showing Transcript levels of *Atf4* (Student’s t-test, N=4). **B** Quantitative analysis showing Transcript levels of *Snap25* (Student’s t-test, N=6). (‘N’ denotes the number of biological replicates) Data are shown as mean ± SEM. (*P < 0.05; **P < 0.01; ***P<0.001; n.s. statistically non-significant).

**Figure S5:**
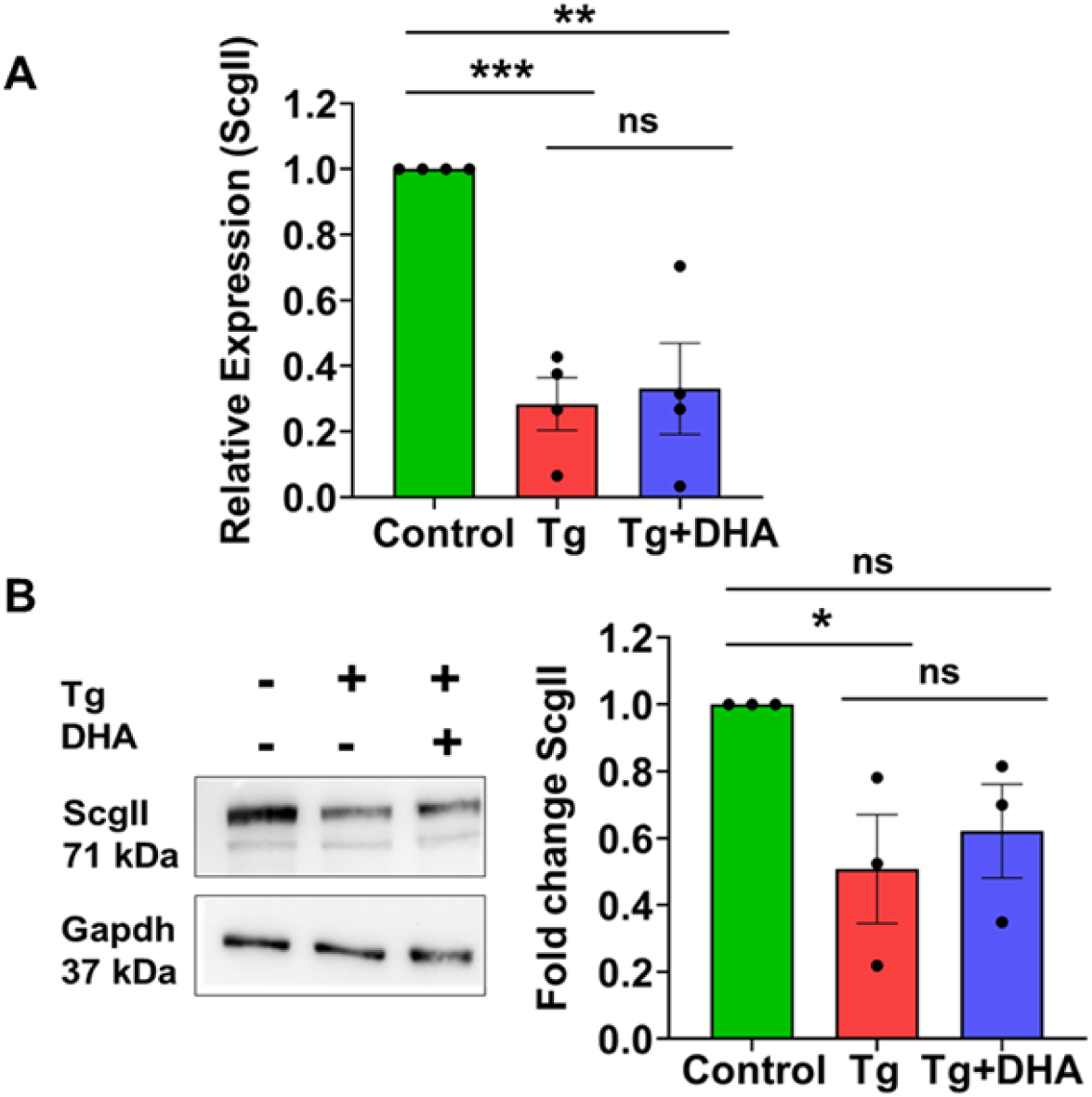
Altered granin expression in ER stressed cells. Quantitative analysis of transcript levels of *ScgII* in Q-RT PCR (Student’s t-test, N=4). **B** Representative Immunoblots of SCGII (left) and its corresponding Quantification (right). GAPDH served as a loading control (Student’s t-test, N=3). (‘N’ denotes the number of biological replicates) Data are shown as mean ± SEM. (*P < 0.05; **P < 0.01; ***P<0.001; n.s. statistically non-significant).

## REFERENCES

Adjibade, P., Grenier St-Sauveur, V., Bergeman, J., Huot, M.-E., Khandjian, E.W., Mazroui, R., 2017. DDX3 regulates endoplasmic reticulum stress-induced ATF4 expression. Sci. Rep. 7, 13832. https://doi.org/10.1038/s41598-017-14262-7

Arora, S., Saarloos, I., Kooistra, R., van de Bospoort, R., Verhage, M., Toonen, R.F., 2017. SNAP-25 gene family members differentially support secretory vesicle fusion. J. Cell Sci. 130, 1877–1889. https://doi.org/10.1242/jcs.201889

Asfari, M., Janjic, D., Meda, P., Li, G., Halban, P.A., Wollheim, C.B., 1992. Establishment of 2-mercaptoethanol-dependent differentiated insulin-secreting cell lines. Endocrinology 130, 167–178. https://doi.org/10.1210/endo.130.1.1370150

Ashraf, G.M., Greig, N.H., Khan, T.A., Hassan, I., Tabrez, S., Shakil, S., Sheikh, I.A., Zaidi, S.K., Wali, M.A., Jabir, N.R., Firoz, C.K., Naeem, A., Alhazza, I.M., Damanhouri, G.A., Kamal, M.A., 2014. Protein misfolding and aggregation in Alzheimer’s disease and Type 2 Diabetes Mellitus. CNS Neurol. Disord. Drug Targets 13, 1280–1293.

Barclay, J.W., Morgan, A., Burgoyne, R.D., 2005. Calcium-dependent regulation of exocytosis. Cell Calcium 38, 343–353. https://doi.org/10.1016/j.ceca.2005.06.012

Barranco, N., Plá, V., Alcolea, D., Sánchez-Domínguez, I., Fischer-Colbrie, R., Ferrer, I., Lleó, A., Aguado, F., 2021. Dense core vesicle markers in CSF and cortical tissues of patients with Alzheimer’s disease. Transl. Neurodegener. 10, 37. https://doi.org/10.1186/s40035-021-00263-0

Beal, M.F., Martin, J.B., 1986. Neuropeptides in neurological disease. Ann. Neurol. 20, 547–565. https://doi.org/10.1002/ana.410200502

Begum, G., Harvey, L., Dixon, C.E., Sun, D., 2013. ER stress and effects of DHA as an ER stress inhibitor. Transl. Stroke Res. 4, 10.1007/s12975-013-0282-1. https://doi.org/10.1007/s12975-013-0282-1

Begum, G., Kintner, D., Liu, Y., Cramer, S.W., Sun, D., 2012a. DHA inhibits ER Ca2+ release and ER stress in astrocytes following in vitro ischemia. J. Neurochem. 120, 622–630. https://doi.org/10.1111/j.1471-4159.2011.07606.x

Begum, G., Kintner, D., Liu, Y., Cramer, S.W., Sun, D., 2012b. DHA inhibits ER Ca2+ release and ER stress in astrocytes following in vitro ischemia. J. Neurochem. 120, 622–630. https://doi.org/10.1111/j.1471-4159.2011.07606.x

Begum, G., Yan, H.Q., Li, L., Singh, A., Dixon, C.E., Sun, D., 2014. Docosahexaenoic Acid Reduces ER Stress and Abnormal Protein Accumulation and Improves Neuronal Function Following Traumatic Brain Injury. J. Neurosci. 34, 3743–3755. https://doi.org/10.1523/JNEUROSCI.2872-13.2014

Bennett, M.C., 2005. The role of alpha-synuclein in neurodegenerative diseases. Pharmacol. Ther. 105, 311–331. https://doi.org/10.1016/j.pharmthera.2004.10.010

Berlier, J.E., Rothe, A., Buller, G., Bradford, J., Gray, D.R., Filanoski, B.J., Telford, W.G., Yue, S., Liu, J., Cheung, C.-Y., Chang, W., Hirsch, J.D., Beechem, J.M., Haugland, Rosaria P., Haugland, Richard P., 2003. Quantitative comparison of long-wavelength Alexa Fluor dyes to Cy dyes: fluorescence of the dyes and their bioconjugates. J. Histochem. Cytochem. Off. J. Histochem. Soc. 51, 1699–1712. https://doi.org/10.1177/002215540305101214

Birinci, Y., Preobraschenski, J., Ganzella, M., Jahn, R., Park, Y., 2020. Isolation of large dense-core vesicles from bovine adrenal medulla for functional studies. Sci. Rep. 10, 7540. https://doi.org/10.1038/s41598-020-64486-3

Brodnanova, M., Hatokova, Z., Evinova, A., Cibulka, M., Racay, P., 2021. Differential impact of imipramine on Thapsigargin- and tunicamycin-induced endoplasmic reticulum stress and mitochondrial dysfunction in neuroblastoma SH-SY5Y cells. Eur. J. Pharmacol. 902, 174073. https://doi.org/10.1016/j.ejphar.2021.174073

Burré, J., Sharma, M., Tsetsenis, T., Buchman, V., Etherton, M.R., Südhof, T.C., 2010. Alpha-synuclein promotes SNARE-complex assembly in vivo and in vitro. Science 329, 1663–1667. https://doi.org/10.1126/science.1195227

Canaff, L., Bevan, S., Wheeler, D.G., Mouland, A.J., Rehfuss, R.P., White, J.H., Hendy, G.N., 1998. Analysis of Molecular Mechanisms Controlling Neuroendocrine Cell Specific Transcription of the Chromogranin A Gene*. Endocrinology 139, 1184–1196. https://doi.org/10.1210/endo.139.3.5851

Cha-Molstad, H., Sung, K.S., Hwang, J., Kim, K.A., Yu, J.E., Yoo, Y.D., Jang, J.M., Han, D.H., Molstad, M., Kim, J.G., Lee, Y.J., Zakrzewska, A., Kim, S.-H., Kim, S.T., Kim, S.Y., Lee, H.G., Soung, N.K., Ahn, J.S., Ciechanover, A., Kim, B.Y., Kwon, Y.T., 2015. N-terminal Arginylation Targets Endoplasmic Reticulum Chaperone BiP to Autophagy Through p62 Binding. Nat. Cell Biol. 17, 917–929. https://doi.org/10.1038/ncb3177

Cicardi, M.E., Marrone, L., Azzouz, M., Trotti, D., 2021. Proteostatic imbalance and protein spreading in amyotrophic lateral sclerosis. EMBO J. 40, e106389. https://doi.org/10.15252/embj.2020106389

Colla, E., Coune, P., Liu, Y., Pletnikova, O., Troncoso, J.C., Iwatsubo, T., Schneider, B.L., Lee, M.K., 2012. Endoplasmic Reticulum Stress Is Important for the Manifestations of α-Synucleinopathy In Vivo. J. Neurosci. 32, 3306–3320. https://doi.org/10.1523/JNEUROSCI.5367-11.2012

Cortès-Saladelafont, E., Tristán-Noguero, A., Artuch, R., Altafaj, X., Bayès, A., García-Cazorla, A., 2016. Diseases of the Synaptic Vesicle: A Potential New Group of Neurometabolic Disorders Affecting Neurotransmission. Semin. Pediatr. Neurol., Update on Neurometabolic Disorders 23, 306–320. https://doi.org/10.1016/j.spen.2016.11.005

de Diego, A.M.G., Ortega-Cruz, D., García, A.G., 2020. Disruption of Exocytosis in Sympathoadrenal Chromaffin Cells from Mouse Models of Neurodegenerative Diseases. Int. J. Mol. Sci. 21, 1946. https://doi.org/10.339/ijms21061946

de Wit, H., A.M. Walter, I. Milosevic, A. Gulyás-Kovács, D. Riedel, J.B. Sørensen, and M. Verhage. 2009. Synaptotagmin-1 Docks Secretory Vesicles to Syntaxin-1/SNAP-25 Acceptor Complexes. Cell. 138:935–946. doi:10.1016/j.cell.2009.07.027.

Dey, S., Savant, S., Teske, B.F., Hatzoglou, M., Calkhoven, C.F., Wek, R.C., 2012. Transcriptional repression of ATF4 gene by CCAAT/enhancer-binding protein β (C/EBPβ) differentially regulates integrated stress response. J. Biol. Chem. 287, 21936–21949. https://doi.org/10.1074/jbc.M112.351783

Dyall, S.C., 2015. Long-chain omega-3 fatty acids and the brain: a review of the independent and shared effects of EPA, DPA and DHA. Front. Aging Neurosci. 7, 52. https://doi.org/10.3389/fnagi.2015.00052

Gagliardi, D., Bresolin, N., Comi, G.P., Corti, S., 2021. Extracellular vesicles and amyotrophic lateral sclerosis: from misfolded protein vehicles to promising clinical biomarkers. Cell. Mol. Life Sci. 78, 561–572. https://doi.org/10.1007/s00018-020-03619-3

Gandasi, N.R., Vestö, K., Helou, M., Yin, P., Saras, J., Barg, S., 2015. Survey of Red Fluorescence Proteins as Markers for Secretory Granule Exocytosis. PLoS ONE 10, e0127801. https://doi.org/10.1371/journal.pone.0127801

Ganten, D., Paul, M., Lang, R.E., 1991. The role of neuropeptides in cardiovascular regulation. Cardiovasc. Drugs Ther. 5, 119–130. https://doi.org/10.1007/BF03029807

Greene, L.A., Tischler, A.S., 1976. Establishment of a noradrenergic clonal line of rat adrenal pheochromocytoma cells which respond to nerve growth factor. Proc. Natl. Acad. Sci. 73, 2424–2428. https://doi.org/10.1073/pnas.73.7.2424

Grindheim, A.K., Hollås, H., Ramirez, J., Saraste, J., Travé, G., Vedeler, A., 2014. Effect of Serine Phosphorylation and Ser25 Phospho-Mimicking Mutations on Nuclear Localisation and Ligand Interactions of Annexin A2. J. Mol. Biol. 426, 2486–2499. https://doi.org/10.1016/j.jmb.2014.04.019

Guha, P., Kaptan, E., Gade, P., Kalvakolanu, D.V., Ahmed, H., 2017. Tunicamycin induced endoplasmic reticulum stress promotes apoptosis of prostate cancer cells by activating mTORC1. Oncotarget 8, 68191–68207. https://doi.org/10.18632/oncotarget.19277

Guo, J., Chang, C., Li, W., 2017. The role of secreted heat shock protein-90 (Hsp90) in wound healing - how could it shape future therapeutics? Expert Rev. Proteomics 14, 665–675. https://doi.org/10.1080/14789450.2017.1355244

Hetz, C., 2012. The unfolded protein response: controlling cell fate decisions under ER stress and beyond. Nat. Rev. Mol. Cell Biol. 13, 89–102. https://doi.org/10.1038/nrm3270

Huang, A.Y., Castle, A.M., Hinton, B.T., Castle, J.D., 2001. Resting (Basal) Secretion of Proteins Is Provided by the Minor Regulated and Constitutive-like Pathways and Not Granule Exocytosis in Parotid Acinar Cells*. J. Biol. Chem. 276, 22296–22306. https://doi.org/10.1074/jbc.M100211200

Hummer, B.H., de Leeuw, N.F., Burns, C., Chen, L., Joens, M.S., Hosford, B., Fitzpatrick, J.A.J., Asensio, C.S., 2017. HID-1 controls formation of large dense core vesicles by influencing cargo sorting and trans-Golgi network acidification. Mol. Biol. Cell 28, 3870–3880. https://doi.org/10.1091/mbc.e17-08-0491

Huttner, W.B., Gerdes, H.H., Rosa, P., 1991. The granin-(chromogranin/secretogranin) family. Trends Biochem. Sci. 16, 27–30. https://doi.org/10.1016/0968-0004(91)90012-K

Huttunen, H.J., Kuja-Panula, J., Rauvala, H., 2002. Receptor for Advanced Glycation End Products (RAGE) Signaling Induces CREB-dependent Chromogranin Expression during Neuronal Differentiation*. J. Biol. Chem. 277, 38635–38646. https://doi.org/10.1074/jbc.M202515200

Jaskulska, A., Janecka, A.E., Gach-Janczak, K., 2020. Thapsigargin-From Traditional Medicine to Anticancer Drug. Int. J. Mol. Sci. 22, 4. https://doi.org/10.3390/ijms22010004

Ji, C., Fan, F., Lou, X., 2017. Vesicle Docking Is a Key Target of Local PI(4,5)P2 Metabolism in the Secretory Pathway of INS-1 Cells. Cell Rep. 20, 1409–1421. https://doi.org/10.1016/j.celrep.2017.07.041

Jin, Y., Zhang, C., Fang, X., Fang, C.-L., Chen, J., Du, R.-L., Hu, Q., Dong, L., Zhu, Z.-Q., Wang, T.-H., 2021. SNAP25 protects primary cortical neurons from hypoxic-ischemic injury associated with CREB signal. Ibrain 7, 1–11. https://doi.org/10.1002/j.2769-2795.2021.tb00058.x

Kim, T., Tao-Cheng, J.-H., Eiden, L.E., Loh, Y.P., 2001. Chromogranin A, an “On/Off” Switch Controlling Dense-Core Secretory Granule Biogenesis. Cell 106, 499–509. https://doi.org/10.1016/S0092-8674(01)00459-7

Kwon, S.E., Chapman, E.R., 2011. Synaptophysin regulates the kinetics of synaptic vesicle endocytosis in central neurons. Neuron 70, 847–854. https://doi.org/10.1016/j.neuron.2011.04.001

Liew, C.W., Bochenski, J., Kawamori, D., Hu, J., Leech, C.A., Wanic, K., Malecki, M., Warram, J.H., Qi, L., Krolewski, A.S., Kulkarni, R.N., 2010. The pseudokinase tribbles homolog 3 interacts with ATF4 to negatively regulate insulin exocytosis in human and mouse β cells. J. Clin. Invest. 120, 2876–2888. https://doi.org/10.1172/JCI36849

Lin, Z., Li, Y., Hang, Y., Wang, C., Liu, B., Li, J., Yin, L., Jiang, X., Du, X., Qiao, Z., Zhu, F., Zhang, Z., Zhang, Q., Zhou, Z., 2022. Tuning the Size of Large Dense-Core Vesicles and Quantal Neurotransmitter Release via Secretogranin II Liquid-Liquid Phase Separation. Adv. Sci. Weinh. Baden-Wurtt. Ger. 9, e2202263. https://doi.org/10.1002/advs.202202263

Lindner, P., Christensen, S.B., Nissen, P., Møller, J.V., Engedal, N., 2020. Cell death induced by the ER stressor thapsigargin involves death receptor 5, a non-autophagic function of MAP1LC3B, and distinct contributions from unfolded protein response components. Cell Commun. Signal. CCS 18, 12. https://doi.org/10.1186/s12964-019-0499-z

Lonze, B.E., Ginty, D.D., 2002. Function and Regulation of CREB Family Transcription Factors in the Nervous System. Neuron 35, 605–623. https://doi.org/10.1016/S0896-6273(02)00828-0

Marciniak, S.J., Chambers, J.E., Ron, D., 2022. Pharmacological targeting of endoplasmic reticulum stress in disease. Nat. Rev. Drug Discov. 21, 115–140. https://doi.org/10.1038/s41573-021-00320-3

McCall, R.B., 1990. Role of neurotransmitters in the central regulation of the cardiovascular system. Prog. Drug Res. Fortschritte Arzneimittelforschung Progres Rech. Pharm. 35, 25–84. https://doi.org/10.1007/978-3-0348-7133-4_2

Mosley, C.A., Taupenot, L., Biswas, N., Taulane, J.P., Olson, N.H., Vaingankar, S.M., Wen, G., Schork, N.J., Ziegler, M.G., Mahata, S.K., O’Connor, D.T., 2007. Biogenesis of the Secretory Granule: Chromogranin A Coiled-Coil Structure Results in Unusual Physical Properties and Suggests a Mechanism for Granule Core Condensation. Biochemistry 46, 10999–11012. https://doi.org/10.1021/bi700704r

Mruk, D.D., Cheng, C.Y., 2011. Enhanced chemiluminescence (ECL) for routine immunoblotting. Spermatogenesis 1, 121–122. https://doi.org/10.4161/spmg.1.2.16606

Neher, E., Sakaba, T., 2008. Multiple roles of calcium ions in the regulation of neurotransmitter release. Neuron 59, 861–872. https://doi.org/10.1016/j.neuron.2008.08.019

Ozcan, L., Tabas, I., 2012. Role of Endoplasmic Reticulum Stress in Metabolic Disease and Other Disorders. Annu. Rev. Med. 63, 317–328. https://doi.org/10.1146/annurev-med-043010-144749

Parchure, A., Tian, M., Stalder, D., Boyer, C.K., Bearrows, S.C., Rohli, K.E., Zhang, J., Rivera-Molina, F., Ramazanov, B.R., Mahata, S.K., Wang, Y., Stephens, S.B., Gershlick, D.C., von Blume, J., 2022. Liquid-liquid phase separation facilitates the biogenesis of secretory storage granules. J. Cell Biol. 221, e202206132. https://doi.org/10.1083/jcb.202206132

Preissler, S., Rato, C., Yan, Y., Perera, L.A., Czako, A., Ron, D., 2020. Calcium depletion challenges endoplasmic reticulum proteostasis by destabilising BiP-substrate complexes. eLife 9, e62601. https://doi.org/10.7554/eLife.62601

Sahu, B.S., Mahata, S., Bandyopadhyay, K., Mahata, M., Avolio, E., Pasqua, T., Sahu, C., Bandyopadhyay, G.K., Bartolomucci, A., Webster, N.J.G., Van Den Bogaart, G., Fischer-Colbrie, R., Corti, A., Eiden, L.E., Mahata, S.K., 2019. Catestatin regulates vesicular quanta through modulation of cholinergic and peptidergic (PACAPergic) stimulation in PC12 cells. Cell Tissue Res. 376, 51–70. https://doi.org/10.1007/s00441-018-2956-1

Sahu, B.S., Obbineni, J.M., Sahu, G., Singh, P.K., Sonawane, P.J., Sasi, B.K., Allu, P.K., Maji, S.K., Bera, A.K., Senapati, S., 2012. Molecular interactions of the physiological anti-hypertensive peptide catestatin with the neuronal nicotinic acetylcholine receptor. J. Cell Sci. 125, 2323–2337.

Salio, C., Lossi, L., Ferrini, F., Merighi, A., 2006. Neuropeptides as synaptic transmitters. Cell Tissue Res. 326, 583–598. https://doi.org/10.1007/s00441-006-0268-3

Satoh, J., Tabunoki, H., Arima, K., 2009. Molecular network analysis suggests aberrant CREB-mediated gene regulation in the Alzheimer disease hippocampus. Dis. Markers 27, 239–252. https://doi.org/10.3233/DMA-2009-0670

Schmittgen, T.D., Livak, K.J., 2008. Analyzing real-time PCR data by the comparative CT method. Nat. Protoc. 3, 1101–1108. https://doi.org/10.1038/nprot.2008.73

Shen, X., Zhang, K., Kaufman, R.J., 2004. The unfolded protein response--a stress signaling pathway of the endoplasmic reticulum. J. Chem. Neuroanat. 28, 79–92. https://doi.org/10.1016/j.jchemneu.2004.02.006

Shi, G., Faúndez, V., Roos, J., Dell’Angelica, E.C., Kelly, R.B., 1998. Neuroendocrine Synaptic Vesicles Are Formed In Vitro by Both Clathrin-dependent and Clathrin-independent Pathways. J. Cell Biol. 143, 947–955.

Smith, S.G., Haynes, K.A., Hegde, A.N., 2020. Degradation of Transcriptional Repressor ATF4 during Long-Term Synaptic Plasticity. Int. J. Mol. Sci. 21, E8543. https://doi.org/10.3390/ijms21228543

Song, W.-J., Yun, J.-H., Jeong, M.-S., Kim, K.-N., Shin, T., Kim, H.-C., Wie, M.-B., 2021. Inhibitors of Lipoxygenase and Cyclooxygenase-2 Attenuate Trimethyltin-Induced Neurotoxicity through Regulating Oxidative Stress and Pro-Inflammatory Cytokines in Human Neuroblastoma SH-SY5Y Cells. Brain Sci. 11, 1116. https://doi.org/10.3390/brainsci11091116

Speese, S., Petrie, M., Schuske, K., Ailion, M., Ann, K., Iwasaki, K., Jorgensen, E.M., Martin, T.F.J., 2007. UNC-31 (CAPS) Is Required for Dense-Core Vesicle But Not Synaptic Vesicle Exocytosis in Caenorhabditis elegans. J. Neurosci. 27, 6150–6162. https://doi.org/10.1523/JNEUROSCI.1466-07.2007

Sulzer, D., Edwards, R.H., 2019. The Physiological Role of α-Synuclein and Its Relationship to Parkinson’s Disease. J. Neurochem. 150, 475–486. https://doi.org/10.1111/jnc.14810

Sweeney, P., Park, H., Baumann, M., Dunlop, J., Frydman, J., Kopito, R., McCampbell, A., Leblanc, G., Venkateswaran, A., Nurmi, A., Hodgson, R., 2017. Protein misfolding in neurodegenerative diseases: implications and strategies. Transl. Neurodegener. 6, 6. https://doi.org/10.1186/s40035-017-0077-5

Szegezdi, E., Reed Herbert, K., Kavanagh, E.T., Samali, A., Gorman, A.M., 2008. Nerve growth factor blocks thapsigargin-induced apoptosis at the level of the mitochondrion viaregulation of Bim. J. Cell. Mol. Med. 12, 2482–2496. https://doi.org/10.1111/j.1582-4934.2008.00268.x

Talbot, S.G., O-charoenrat, P., Sarkaria, I.S., Ghossein, R., Reddy, P., Ngai, I., Cordeiro, C.N., Wong, R.J., Kris, M.G., Rusch, V.W., Singh, B., 2004. Squamous cell carcinoma related oncogene regulates angiogenesis through vascular endothelial growth factor-A. Ann. Surg. Oncol. 11, 530–534. https://doi.org/10.1245/ASO.2004.03.014

Teleanu, R.I., Niculescu, A.-G., Roza, E., Vladâcenco, O., Grumezescu, A.M., Teleanu, D.M., 2022. Neurotransmitters-Key Factors in Neurological and Neurodegenerative Disorders of the Central Nervous System. Int. J. Mol. Sci. 23, 5954. https://doi.org/10.3390/ijms23115954

Thomas-Reetz, A.C., De Camilli, P., 1994. A role for synaptic vesicles in non-neuronal cells: clues from pancreatic β cells and from chromaffin cells. FASEB J. 8, 209–216. https://doi.org/10.1096/fasebj.8.2.7907072

Tsuboi, T., Fukuda, M., 2005. The C2B Domain of Rabphilin Directly Interacts with SNAP-25 and Regulates the Docking Step of Dense Core Vesicle Exocytosis in PC12 Cells *.J. Biol. Chem. 280, 39253–39259. https://doi.org/10.1074/jbc.M507173200

Ulrich, G., Ciesielski-Treska, J., Taupenot, L., Bader, M.-F., 2002. Chromogranin A-activated microglial cells induce neuronal apoptosis. Ann. N. Y. Acad. Sci. 971, 560–562. https://doi.org/10.1111/j.1749-6632.2002.tb04527.x

van Meerloo, J., Kaspers, G.J.L., Cloos, J., 2011. Cell Sensitivity Assays: The MTT Assay, in: Cree, I.A. (Ed.), Cancer Cell Culture: Methods and Protocols, Methods in Molecular Biology. Humana Press, Totowa, NJ, pp. 237–245. https://doi.org/10.1007/978-1-61779-080-5_20

Walter, A.M., Wiederhold, K., Bruns, D., Fasshauer, D., Sørensen, J.B., 2010. Synaptobrevin N-terminally bound to syntaxin–SNAP-25 defines the primed vesicle state in regulated exocytosis. J. Cell Biol. 188, 401–413. https://doi.org/10.1083/jcb.200907018

Wang, B., Zhang, J., Liu, X., Chai, Q., Lu, X., Yao, X., Yang, Z., Sun, L., Johnson, S.F., Schwartz, R.C., Zheng, Y.-H., 2022. Protein disulfide isomerases (PDIs) negatively regulate ebolavirus structural glycoprotein expression in the endoplasmic reticulum (ER) via the autophagy-lysosomal pathway. Autophagy 1–18. https://doi.org/10.1080/15548627.2022.2031381

Wang, J., Xu, W., Kong, Y., Huang, J., Ding, Z., Deng, M., Guo, Q., Zou, W., 2019. SNAP-25 Contributes to Neuropathic Pain by Regulation of VGLuT2 Expression in Rats. Neuroscience 423, 86–97. https://doi.org/10.1016/j.neuroscience.2019.10.007

Wang, X., Huang, T., Bu, G., Xu, H., 2014. Dysregulation of protein trafficking in neurodegeneration. Mol. Neurodegener. 9, 31. https://doi.org/10.1186/1750-1326-9-31

Wemhöner, A., Frick, M., Dietl, P., Jennings, P., Haller, T., 2006. A Fluorescent Microplate Assay for Exocytosis in Alveolar Type II Cells. J. Biomol. Screen. 11, 286–95. https://doi.org/10.1177/1087057105285284

Westerink, R.H.S., Ewing, A.G., 2008. The PC12 cell as model for neurosecretion. Acta Physiol. Oxf. Engl. 192, 273–285. https://doi.org/10.1111/j.1748-1716.2007.01805.x

Wilkinson, B., Gilbert, H.F., 2004. Protein disulfide isomerase. Biochim. Biophys. Acta 1699, 35–44. https://doi.org/10.1016/j.bbapap.2004.02.017

Yang, X., Lou, J., Shan, W., Ding, J., Jin, Z., Hu, Y., Du, Q., Liao, Q., Xie, R., Xu, J., 2021. Pathophysiologic Role of Neurotransmitters in Digestive Diseases. Front. Physiol. 12.

Yoshida, H., 2007. ER stress and diseases. FEBS J. 274, 630–658. https://doi.org/10.1111/j.1742-4658.2007.05639.x

Zhang, Z., Wu, Y., Wang, Z., Dunning, F.M., Rehfuss, J., Ramanan, D., Chapman, E.R., Jackson, M.B., 2011. Release mode of large and small dense-core vesicles specified by different synaptotagmin isoforms in PC12 cells. Mol. Biol. Cell 22, 2324–2336. https://doi.org/10.1091/mbc.E11-02-0159

Zorec, R., 1996. Calcium signaling and secretion in pituitary cells. Trends Endocrinol. Metab. TEM 7, 384–388. https://doi.org/10.1016/s1043-2760(96)00169-5

Zou, L., Tian, Y., Zhang, Z., 2021. Dysfunction of Synaptic Vesicle Endocytosis in Parkinson’s Disease. Front. Integr. Neurosci. 15.

